# Preference and familiarity mediate spatial responses of a large herbivore to experimental manipulation of resource availability

**DOI:** 10.1101/757757

**Authors:** Nathan Ranc, Paul R. Moorcroft, K. Whitney Hansen, Federico Ossi, Tobia Sforna, Enrico Ferraro, Alessandro Brugnoli, Francesca Cagnacci

## Abstract

The link between spatio-temporal resource patterns and movement behaviour is a key ecological process, however, limited experimental support has been produced at the home range scale. In this study, we analysed the spatial responses of a resident large herbivore (roe deer *Capreolus capreolus*) during an *in situ* manipulation of a concentrated food resource. Specifically, we experimentally altered feeding site accessibility to roe deer and recorded (for 25 animal-years) individual responses by GPS tracking. We found that, following the loss of their preferred resource, roe deer actively tracked resource dynamics leading to more exploratory movements, and larger, spatially-shifted home ranges. Then, we showed, for the first time experimentally, the importance of site fidelity in the maintenance of large mammal home ranges by demonstrating the return of individuals to their familiar, preferred resource despite the presence of alternate, equally-valuable food resources. This behaviour was modulated at the individual level, where roe deer characterised by a high preference for feeding sites exhibited more pronounced behavioural adjustments during the manipulation. Together, our results establish the connections between movement, space-use, individual preference, and the spatio-temporal pattern of resources in home ranging behaviour.

## Introduction

Animals move to change the environmental context they experience^1^, including abiotic conditions, the presence of predators and competitors, and the availability of resources. Because foraging efficiency can be linked to individual fitness^2^, food acquisition is thought to be a primary driver underlying animal movements^3^. Consequently, space-use represents the geographic realization of optimizing fitness as a function of resource availability and acquisition costs^4^.

Food resources are usually dynamic in both space and time^5^. In the case of herbivores, animals typically feed on vegetation distributed in patches, which are characterized by important temporal variations in quantity and quality^6^. In this context, strong spatio-temporal gradients in resource availability at either landscape or regional scales appear to drive migration and nomadism tactics^7^. In many herbivore populations, however, individuals show a high year-round fidelity to a spatially-localized home range. It has been suggested that the foraging benefits of site familiarity, where resources are constant or predictable, are responsible for the formation of a stable home range (see ^8^ for a review). While the home range has traditionally been perceived as a relatively static space-use tactic, recent evidence suggests that animals have sub-seasonal home ranges^9^ i.e., focus their movements into particular areas in response to seasonal variation in local resource availability. For example, sub-seasonal home ranges are a ubiquitous behavioural tactic across a wide ecological gradient in roe deer (*Capreolus capreolus*)^10^.

The link between movement behaviour and resource dynamics is less clear when observing home ranging behaviour than migration or nomadism^3^ because of the difficulty to quantify spatio-temporal variability in resource heterogeneity at small spatial scales^10^. In this study, we address this issue by experimentally manipulating the spatio-temporal patterns of food availability within home ranges. *In situ* food manipulation experiments have a long history in the study of population dynamics, with a primary focus on understanding the numerical response to food supplementation^11^, and of animal communities^12^. Although these field experiments have provided fundamental insights in animal ecology, they have seldom been combined with the emerging technological capabilities of animal tracking^13^ to investigate the implications of food availability on individual movements and space-use at larger spatial scales. In a rare example of field experiment in large herbivores, white-tailed deer (*Odocoileus virginianus*) shifted their core-area i.e., familiar areas of use, in response to novel food supplementation^14^. In turn, great tits (*Parus major*) showed personality-dependent variability in responses to an alteration of resource distribution^15^.

Our research builds upon these two studies by investigating the spatial responses of a large herbivore, roe deer, to an experimental *in situ* manipulation of a high-quality, concentrated food resource in relation to both individual resource preferences and site familiarity. Roe deer, as solitary browsers with limited fat reserves^16^, exhibit a tight association between movement and resource dynamics^17^ with a strong plasticity to adapt its resource acquisition at different spatio-temporal scales^18–20^. In contrast to group-living ungulates, their foraging decisions are expected to be clearly expressed at the level of individuals.

We tagged roe deer in the Eastern Italian Alps with GPS units and followed their movements during transitory alterations of food availability at supplemental feeding sites (FS) i.e., discrete resource patches with an identifiable resource value distinguishable from the vegetation matrix^4,5^. The six-week experiment consisted of three two-week phases – pre-closure, closure and post-closure. During the closure phase, we physically restricted the accessibility of food at the most attended feeding site of each individual (hereafter referred to as Manipulated, M, and considered as most familiar; Fig. 1). Throughout the experiment, roe deer had access to two alternative resources within the broader landscape: alternate feeding sites (A), whose food provisioning was held constant, and natural vegetation (V). The experiment was conducted during winter, when food scarcity limits roe deer foraging performance, and individuals are most inclined to adjust their spatial behaviour to continue meeting their energy requirements^21^. The experiment therefore mimics – on free-ranging animals – the variation in the availability of concentrated, high-reward resources akin to watering holes for savannah ungulates^22^ and feral horses (*Equus ferus*)^23^, or fruit trees for hornbills (C*eratogymna atrata* and *C. cylindricus*)^24^ and frugivorous primates^25,26^.

**Figure 1.**
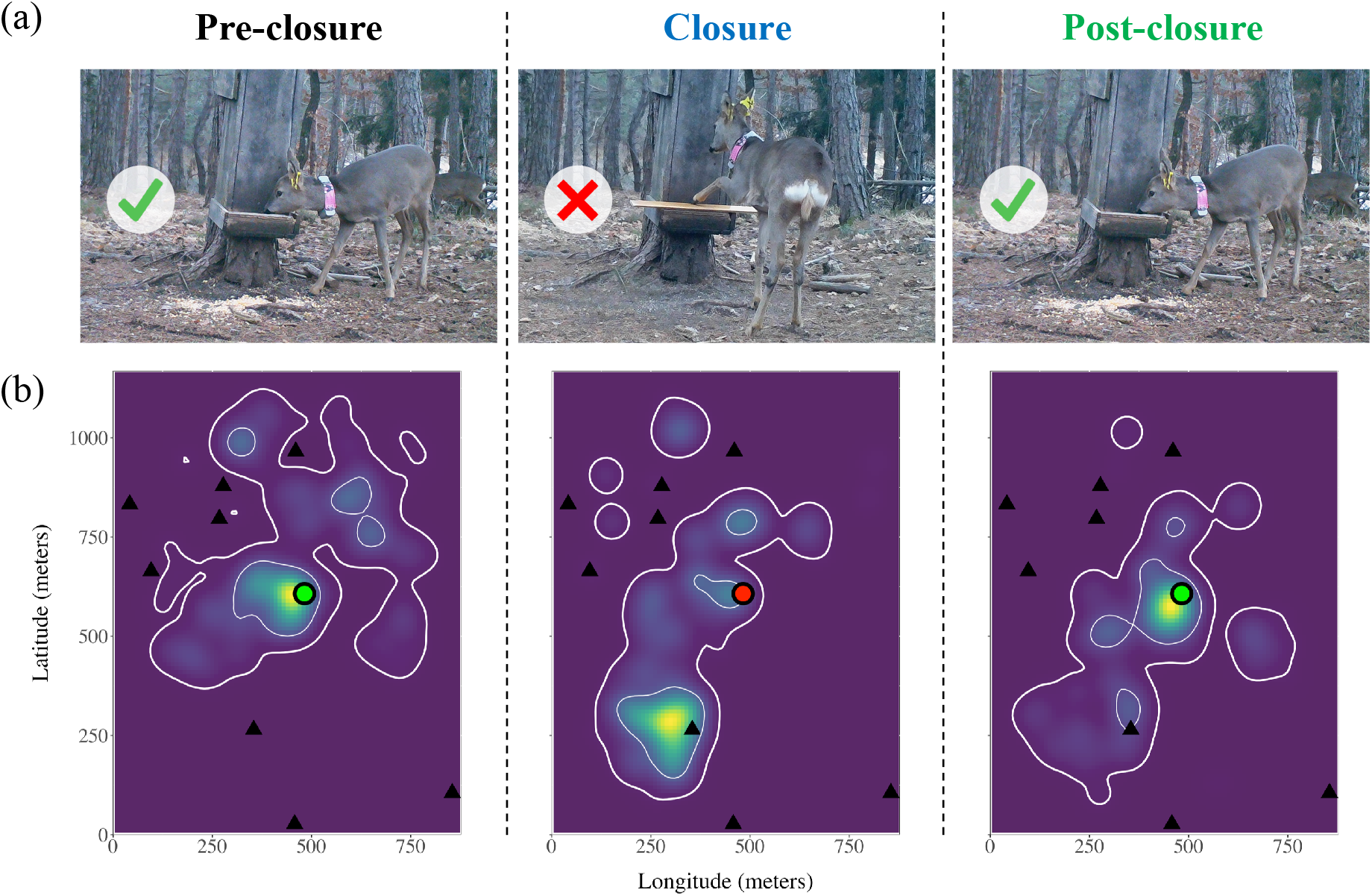
Schematic representation of the experiment. (a) The manipulation consists of a transitory alteration of resource accessibility at a manipulated (M) feeding site. (b) The experiment is expected to lead to spatial responses in the monitored roe deer, and in particular in a shift of use from M (green/red dot, change of colour denoting the alteration of accessibility) towards alternative resources – alternate feeding sites (A; black triangles) or the natural vegetation (V; underlying matrix). In particular, this can lead to spatio-temporal dynamics in space-use (utilization distribution: colour gradient; 95% and 50% contour lines: thick and thin white lines, respectively; data from roe deer F5-2018).

Our initial hypothesis states that individuals alter their movement behaviours and consequently space-use patterns to track dynamics in resource availability (H1; Table 1). We predicted that the loss of a key foraging resource should lead to larger (P1.1) and spatially-shifted (P1.2) home ranges, resulting from more explorative movements (P1.3). Furthermore, we predicted that roe deer reduced the intensity of use of the manipulated, familiar FS (M) when food accessibility was prevented (P1.4a) and compensated for this loss by using alternate, accessible FS (A; P1.4b).

**Table 1.**
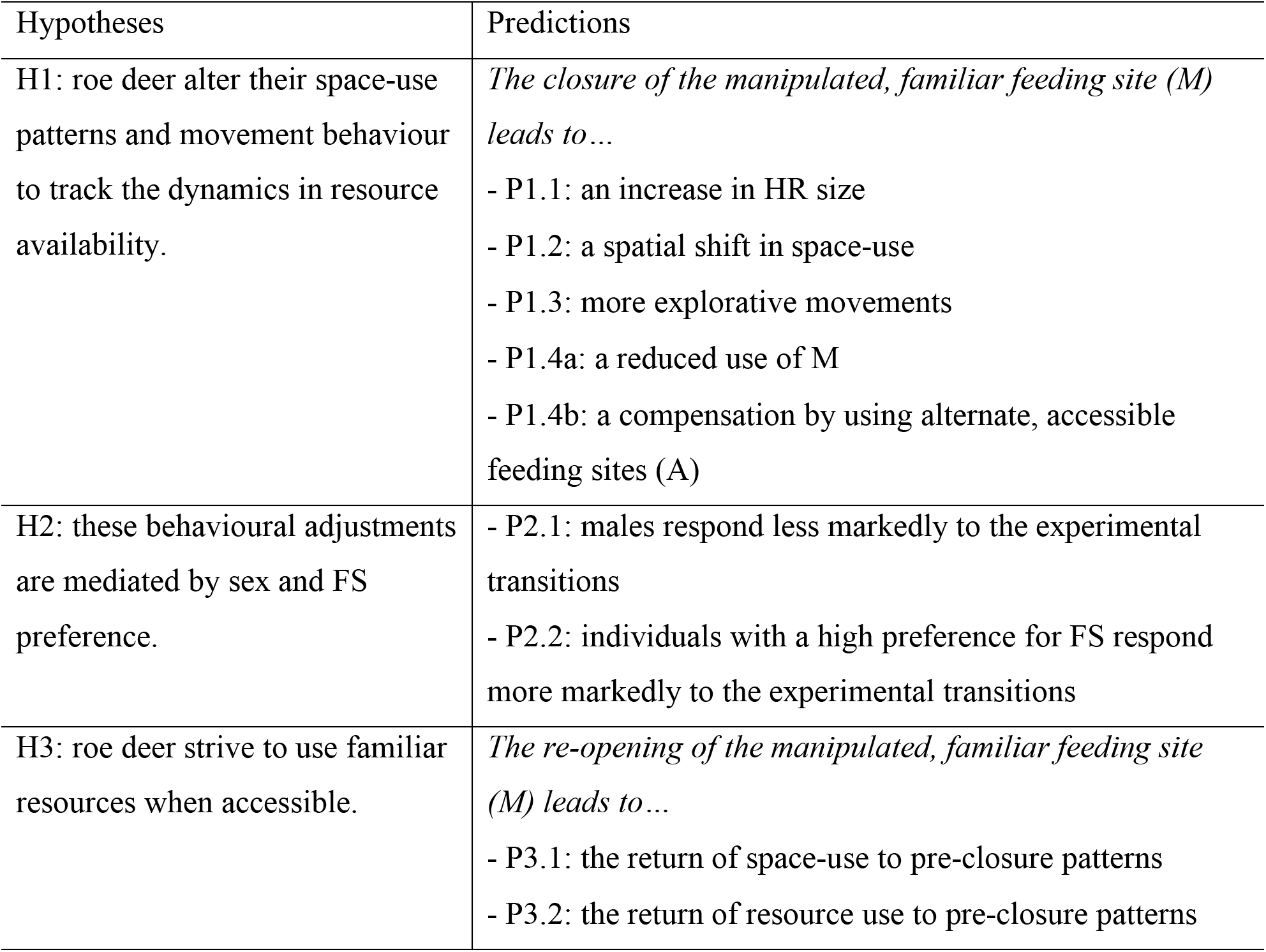
Hypotheses and corresponding predictions.

| Hypotheses | Predictions |
| --- | --- |
| H1: roe deer alter their space-use patterns and movement behaviour to track the dynamics in resource availability. | *The closure of the manipulated, familiar feeding site (M) leads to...*<br>- P1.1: an increase in HR size<br>- P1.2: a spatial shift in space-use<br>- P1.3: more explorative movements<br>- P1.4a: a reduced use of M<br>- P1.4b: a compensation by using alternate, accessible feeding sites (A) |
| H2: these behavioural adjustments are mediated by sex and FS preference. | - P2.1: males respond less markedly to the experimental transitions<br>- P2.2: individuals with a high preference for FS respond more markedly to the experimental transitions |
| H3: roe deer strive to use familiar resources when accessible. | *The re-opening of the manipulated, familiar feeding site (M) leads to...*<br>- P3.1: the return of space-use to pre-closure patterns<br>- P3.2: the return of resource use to pre-closure patterns |

We further hypothesized that the behavioural adjustments to changes in resource availability would vary between individuals (H2; Table 1). In particular, because roe deer males have been shown to maintain a high year-round fidelity to their summer territory^27^, we predicted that they would respond less markedly to the experiment than females (P2.1). We also predicted the responsiveness of roe deer to be positively influenced by the individual’s prior preference for FS (P2.2).

If the spatial patterns of roe deer home ranges result from the benefits of site familiarity^28,29^, animals should strive to use familiar areas and resources when accessible (H3; Table 1). Accordingly, we predicted that when initial conditions of food accessibility are re-established after perturbation, the initial space-use patterns would be restored (P3.1), following a return to high use of the familiar FS (M; P3.2).

## Results

### Space-use and movement responses to alteration of resource availability

Roe deer space-use changed significantly during the experiment: the size of both home ranges (95% UD isopleth; Fig. 2a; Supplementary S4: Table S1) and core areas (50% isopleth; Fig. 2b; Supplementary S4: Table S2) increased significantly during the experimental closure (P1.1). On average, home range size increased from 27.99 ha (*σ*=11.02) during pre-closure to 34.97 ha (*σ*=10.17) during closure, and settled to 29.40 ha (*σ*=9.27) during post-closure. Core area size followed a similar trend with averages of 4.23 ha (*σ*=2.34), 5.85 ha (*σ*=2.33) and 4.98 ha (*σ*=2.09), respectively.

**Figure 2.**
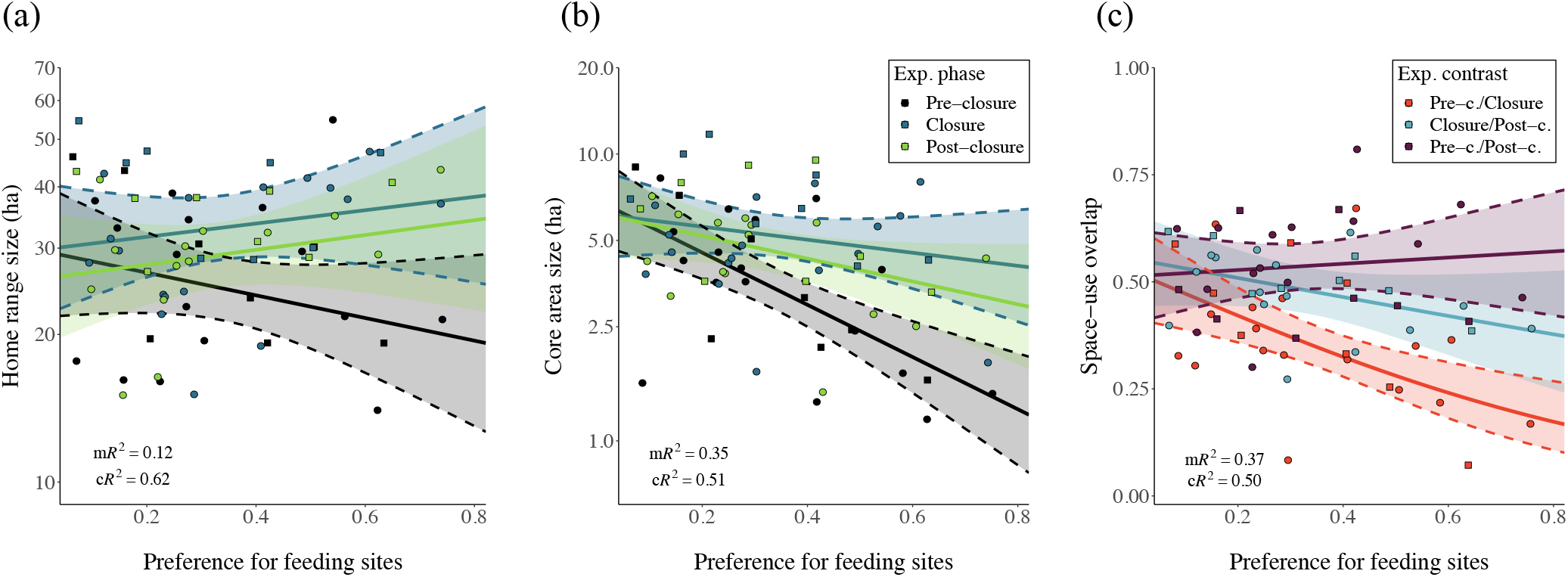
Changes in roe deer space-use patterns – home range size (y-axis, panel a), core area size (y-axis, panel b) and space-use overlap (y-axis, panel c) – as a function of preference for feeding sites (x-axis) and experimental phase (colour; panels a and b) and phase contrast (colour; panel c). Observations are represented as dots (females) and squares (males) with a slight jitter, and model predictions as solid lines (95% confidence intervals: ribbons). Parameter estimates and associated significance can be found in Supplementary S4: Tables S1, S2 and S5.

Home range and core area sizes were influenced by individual preference for FS (*h*_*FS*_) and there was an interaction between *h*_*FS*_ and experimental phase: individuals with a high *h*_*FS*_ tended to have smaller home ranges during the pre-closure, smaller core areas overall, and stronger increases in both home range and core area sizes following the experimental closure (Fig. 2a-b; Supplementary S4: Tables S1, S2; P2.2). There was no significant effect of sex or interactions between sex and experimental phase on home range size (Supplementary S4: Table S3; P2.1), but a marginally significant interaction between sex and experimental phase on core area size (Supplementary S4: Table S4), with responses to closure tending to be slightly larger for males. Overall, the models quantifying the changes in observed home range and core area sizes accounted for a high proportion of the total variance (conditional coefficient of determination, cR^2^: 0.62 and 0.51, respectively).

Alongside home range size, the spatial pattern of roe deer home ranges shifted dramatically following the experimental closure (Fig. 2c): the degree of space-use overlap between pre-closure and closure phases was significantly lower (mean=0.370, CI=0.301-0.405; P1.2) than the overlap between the temporally-separated pre- and post-closure phases (mean=0.535, CI=0.475-0.594; P3.1). Space-use overlap was significantly affected by *h*_*FS*_ (Fig. 2c; P2.2), with higher *h*_*FS*_ being associated to larger space-use shifts following closure (Supplementary S4: Table S5). However, there was no apparent influence of sex in the observed space-use patterns (Supplementary S4: Table S6; P2.1). The model predicting space-use overlap accounted for an important proportion of the variance (cR^2^=0.50).

Underpinning these changes in home range size and space-use patterns were important changes in roe deer movement behaviour during the experiment. Average hourly step length during the pre-closure phase was 60.32 m (*σ*=85.79); during closure it increased to 74.26 m (*σ*=108.11); and during post-closure it decreased to 68.18 m (*σ*=96.61, P1.3). In general, males (Supplementary S5: Fig. S1, right-hand panels, Table S1; P2.1), and individuals associated with high *h*_*FS*_ values (Supplementary S5: Fig. S1, top panels, Table S1; P2.2) were characterized by stronger increases in step length during the closure phase. In addition, roe deer movements were more persistent during the closure phase, as shown by a significant decrease in the mean absolute turning angle for males with a high *h*_*FS*_ (Supplementary S5: Fig. S2, top-right panel, Tables S2, S3; P1.3, P2.1, P2.2).

### Resource use responses to alteration of availability

The spatio-temporal dynamics of resources availability during the experiment led to important shifts in resource use (Fig. 3; Supplementary S6: Table S1). On average, the proportion of use of the manipulated FS (M) dropped from 31% during the pre-closure phase to 4% during closure (P1.4a), and then rebounded to 19% in the post-closure phase (P3.2). This decrease in the use of M during the closure phase was partially compensated by elevated use of the alternate FS (A) – which increased from 3% to 16% following closure (P1.4b), and an increase of the use of vegetation (V) from 66% to 80% following closure. During the post closure, use of A and V declined to 9% and 72%, respectively. The shifts in resource use were very consistent among animal-years for M and A but were more variable for V (Fig. 3, top panels).

**Figure 3.**
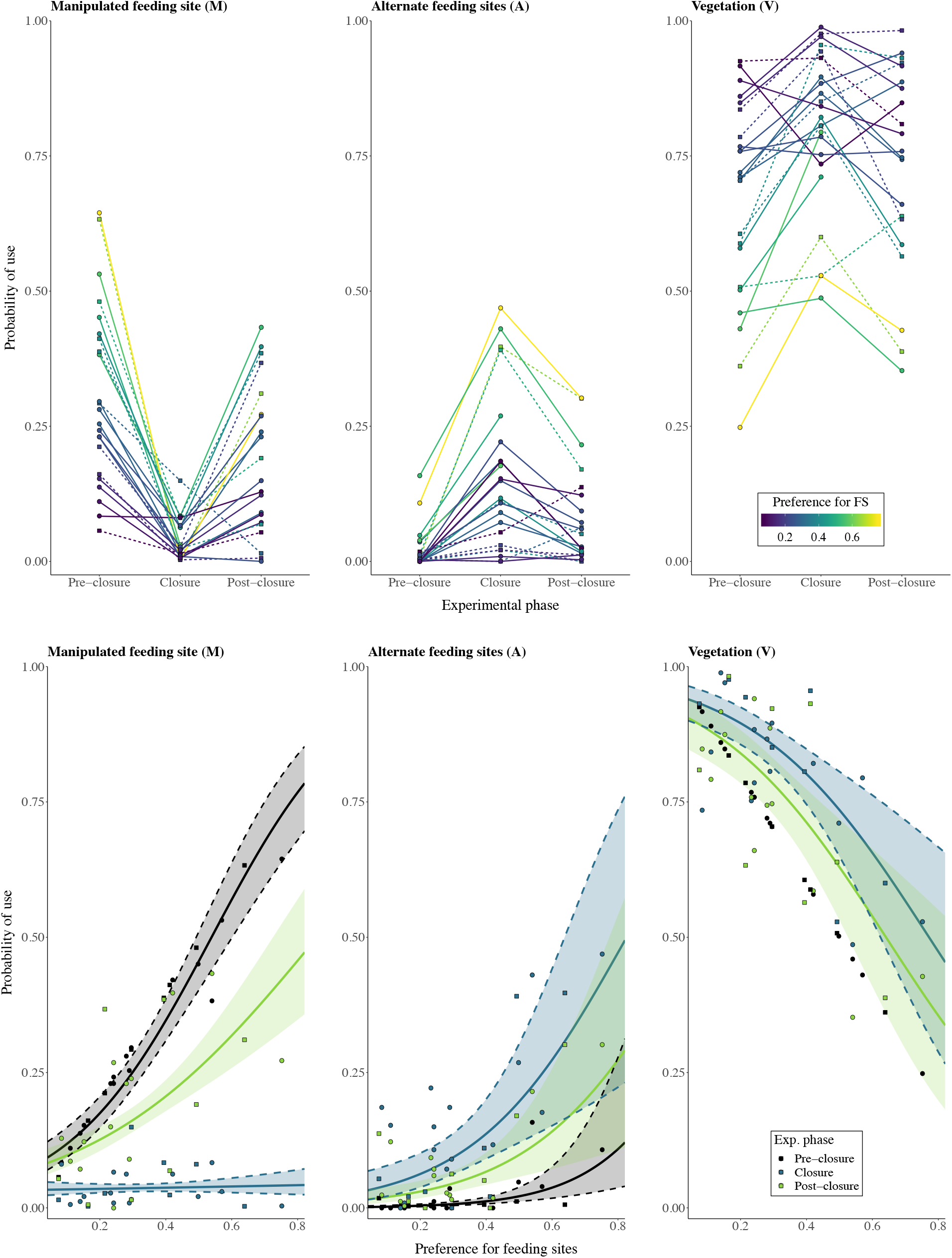
Roe deer shifts in resource use during the experiment – manipulated feeding site (M, left), alternate feeding sites (A, centre) and vegetation (V, right). Top panel: mean proportional use (dots and lines) as a function of the experiment phase (x-axis) and preference for feeding sites (colour scale). Bottom panel: predictions of the resource use models (*u*_*M*,*t*_, *u*_*A*,*t*_ and *u*_*V*,*t*_; estimate: solid lines; 95% confidence interval: ribbon) and mean relative use (females: dots; males: squares) as a function of the experiment phase (colour) and preference for feeding sites (x-axis). The model predictions do not consider resource lags at 1, 2 and 24 h nor the influence of *Sex* (although selected in the final model for *u*_*A*,*t*_) for clarity and conciseness. Parameter estimates and associated significance can be found in Supplementary S6: Table S1.

Roe deer preference for FS significantly influenced how animals used the three resource types and, in particular, interacted with experimental phase for M and A (Fig. 3, bottom panels; Supplementary S6: Table S1). Roe deer characterized by a high *h*_*FS*_ had significantly higher use of M during pre-closure (by definition) and post-closure, as well as consistently lower use of V. High *h*_*FS*_ animals were associated with stronger decreases in use of M and larger increases in the use of A during closure (P2.2). This compensation for A during closure was stronger for females (Supplementary S6: Fig. S1, Table S1; P2.1). However, sex did not influence the use of M or V (Supplementary S6: Table S2; P2.1). Overall, the fitted models accounted for a high proportion of the variance in resource use (cR^2^: 0.35, 0.21 and 0.31 for M, A and V, respectively).

## Discussion

The results of this field resource manipulation experiment provide direct evidence for the tight coupling between the spatio-temporal distribution of resources and consequently spatially-restricted movements of a resident large herbivore. Specifically, we show that roe deer track resource dynamics (Fig. 3; H1), which leads to changes in their space-use (Fig. 2) and underpinning movements (Supplementary S5: Figs S1, S2), and that individual traits, especially resource preference, mediate these behavioural adjustments (H2). In addition, we show that roe deer exhibit a high attraction to familiar locations, a process which leads to site fidelity (H3). As far as we are aware, this is the first experimental demonstration of these interdependencies in a large mammalian herbivore.

The experimental alterations of food availability led to larger, spatially-shifted home ranges (Fig. 3), and more explorative movements by roe deer (Supplementary S5: Figs S1, S2), thereby directly establishing the connections between movement, space-use and the spatio-temporal patterns of resources.

In a previous observational study, elk were shown to alternate between two movement modes: a low speed and high sinuosity mode thought to be within-patch area-restricted search, and a high speed and low sinuosity mode between resource patches^38^. In our experimental study, we can directly link these movement modes to changing resources: the exploratory movements of roe deer (high velocity and low sinuosity) observed during the closure phase (P1.3) suggested that the animals were motivated to find alternate resource patches when their familiar feeding site (FS) became inaccessible, thereby increasing (P1.1), but mainly shifting (P1.2), their home range. While changes in home range size and location following resource manipulation have been found in studies of lizards^39^, birds^15^ and voles^40^, to date, there have been few experimental investigations of the connections between space-use and the spatio-temporal distribution of resources in large mammals.

In an earlier study, white-tailed deer shifted their home range core towards the vicinity of newly deployed FS^14^. Our study builds upon these results by demonstrating multiple, successive responses to resource manipulation, linking measured changes in underlying fine-scale movement behaviour of individuals to resulting patterns of space-use that indicate dynamic resource tracking behaviour by roe deer (H1). Although roe deer increased their use of the vegetation matrix during the closure phase, individuals compensated the loss of their manipulated FS (M), to a large degree, by shifts in their movements and space-use towards alternate FS (A; Figs 3; P1.4a-b). Consequently, individuals maintained a high overall use of FS throughout the experiment.

While resource tracking behaviour may be expected under the optimal foraging theory, other individual- and population-based factors, such as social fences due to territoriality or density-dependent resource competition, can constrain the movement responses of individuals to changes in the spatial distribution of resources^41^. The marked responses shown here are likely the result of two non-mutually exclusive conditions. First, with the exception of adult males during spring and summer^42^, roe deer do not generally defend territories, and consequently their spatial distribution can approximate that of an ideal free distribution^43^. In fact, territorial tenure^27^ may explain the marginally different response of males (P2.1), specifically their tendency to have larger core areas (Supplementary S4: Table S4), more explorative movements (Supplementary S5: Tables S1, S2) and lower resource compensation than females following closure (Supplementary S6: Table S1, Fig. S1). Second, while intra- and inter-specific competition in herbivores is largely linked to resource depletion^6^, this density-dependent constraint of food availability was prevented by providing *ad libitum* forage at the FS.

This study demonstrates that inter-individual variation in preference for FS strongly mediated the responses of roe deer movement patterns, space-use and resource use to changes in the spatio-temporal distribution of resources (H2). During the closure phase, the changes in all measured variables were of larger magnitude for individuals associated with a high FS preference (P2.1). The influence of FS preference was particularly striking in the shifts of space-use (Fig. 2c) and in the compensating use of alternate FS following the loss of the familiar resource (Fig. 3). The observed inter-individual differences in FS preference (Supplementary S3: Table S1) may be linked to either the environment the individuals were exposed to, or a property of the individuals themselves. In our experimental setting, all roe deer had access to at least one FS provided with *ad libitum* food where use was not prevented by inter-individual competition^17^. Hence, we could quantitatively approximate FS preference as the relative use of FS over natural vegetation. Preference for FS was derived from the relative use of FS, and so our definition implies a conditionality on both the physiological state of the individual and on the relative nutritional value of the provisioned food (corn) over natural vegetation in a given season. In addition, the dynamics in the quality and quantity of natural browse – either spatial (e.g., between home ranges) and/or temporal (e.g., between years) – could lead to temporal variations in FS preference. Indeed, FS preference was higher in 2017 than in 2018 for all roe deer manipulated in these consecutive years (Supplementary S3: Table S1). Preference can therefore be considered a dynamic variable^30^ that we evaluated at the individual level over a short period of relative stability (pre-closure phase in each winter). We considered the temporal extent of our experiment (*ca* 6 weeks) short enough to consider FS preference for each animal-year to be relatively constant, because in this time period the physiological conditions and vegetation nutritional value would not vary substantially or consistently.

Animals attending FS benefit from exploiting a forage-rich location but may risk elevated intra- and inter-specific contacts, anthropogenic disturbance, and an increased susceptibility to predation by humans. In roe deer, individual behavioural profile (e.g., body temperature at capture) correlates with the use of risky but profitable habitats such as open areas, suggesting that variations in personality could lead to individual differences in habitat use^44^. By analogy, FS preference could be associated to bold or risk-taking personalities. Interestingly, preference for FS tended to correlate with individual body temperature at capture (Pearson’s r = − 0.37 p-value = 0.084; Supplementary S8), with bolder individuals (lower temperature) using FS more intensely. Personality may not only lead individuals to use resources to a different extent, but also condition their tendency to track spatio-temporal resource dynamics^45^. For instance, risk-taking personalities tend to be associated with explorations of novel environments^46^, as shown experimentally in great tits following the loss of a familiar foraging area^15^.

During the post-closure phase of the experiment, roe deer increased their use of familiar FS (M), whose food accessibility had been restored after a transitory restriction (Fig. 3, left-hand panels; P3.1), and home ranges shifted back to pre-closure patterns, as suggested by the high spatial overlap between temporally-disjointed pre- and post-closure space-use (Fig. 2c; P3.2). The restoration of these pre-manipulation patterns supports the hypothesis that site familiarity provides inherent benefits to animals maintaining a home range (H3)^47^. These results are coherent with published literature demonstrating that ungulates tend to select for previously visited locations i.e., site familiarity^48,49^.

Unlike most observational studies, our experimental approach allowed us to contrast two concentrated resources (M and A) of equal nutritional value but of distinct familiarity, and hence to separate the effects of resource tracking from those of familiarity. Resource tracking can explain roe deer use of alternate FS (A) to compensate the inaccessibility of the familiar, manipulated FS (M) during closure. However, the systematic return of roe deer to M during the post-closure phase, while both alternative resources were accessible, cannot be explained by resource tracking alone. Our experiment therefore suggests that roe deer were actively selecting for familiar areas and that site familiarity has an inherent value.

In observational studies of animal movement, a spurious familiarity effect can occur when an important factor influencing animal behaviour is not considered, and the re-visitation of particular locations is interpreted as an evidence for site familiarity selection^50^. However, this confounding effect is unlikely to affect the results of this experiment. First, corn was delivered *ad libitum* across all FS (M or A) i.e., homogeneous foraging value. Second, the FS were located in comparable environments and especially in relation to cover, a factor that largely influences roe deer movements and space-use^51,52^. Third, and most importantly, the specific identities of M and A varied interchangeably between individuals. Hence, we conclude that the return to pre-closure patterns of foraging behaviour and space-use are unlikely to be result of variations in the characteristics of specific FS, but rather of an inherent familiarity effect. In roe deer, site familiarity could allow a more profitable exploitation of natural forage, as seen in bison (*Bison bison)*^49^ or reduce intraspecific competition for such resource (see ^28^ for a theoretical argument). Alternatively, the attraction to familiar areas could be related to a predator (natural or human) avoidance tactic^53^.

Our results imply that when resource patterns are changing, individual behavioural decisions probably reflect a trade-off between the advantages of site familiarity and resource tracking. Species with little fat reserves need a constant, high-nutritional intake, and may hence be required to rapidly adjust their movements away from their most familiar areas to track spatio-temporal resource dynamics, as seen in roe deer during the closure phase of this experiment or in great tits^15^. It would be interesting to investigate whether capital breeding species, with greater capacity to buffer transitory shortages of food availability, are more reluctant to abandon familiar locations.

Ultimately, site familiarity is the manifestation of an animal’s ability to acquire spatial information, in particular by means of spatial memory^8^. Large herbivores are capable of memorizing the location and profitability of resources^49^. In this study, it is likely that the variations in roe deer responses to resource changes that are not explained by preference may be the result of individual prior experience and knowledge of the status and distribution of concentrated resources. An interesting avenue for further studies will be to evaluate the role of these cognitive processes on individual foraging decisions.

## Methods

### Study area

Roe deer is the primary large herbivore in the study area (7-8 individuals km^−2^), located in the north-eastern Italian Alps (Argentario range, Autonomous Province of Trento). The area is characterized by a continental climate (mean temperature of January: 1.0 °C; July: 21.0 °C; mean annual rainfall: 966 m) with occasional snow cover, and is largely forested (80%). It encompasses four hunting reserves in which selective hunting occurs between September and December (see Supplementary S1 for further details).

Supplemental feeding management of roe deer is conducted year-round at > 50 distinct feeding sites (FS; Supplementary S2: Fig. S1). FS are typically shaped as wooden hopper dispensers that provide a continuous supply of corn accessible through a tray (Fig. 1a). They are managed by private hunters for roe deer but are also attended sporadically by red deer (*Cervus elaphus*).

### Experimental design

We took advantage of roe deer use of a focal, identifiable resource – the FS – to design an *in situ* experimental manipulation of resource availability. We created three successive experimental phases of the availability of this resource – pre-closure, closure and post-closure – by physically managing the accessibility of food at the FS. During the closure phase, access to forage at FS was transitorily restricted by placing wooden boards obstructing the tray; these were then removed again in the post-closure phase (Fig. 1).

The experiment was conducted between January and April, when roe deer use of supplemental feeding is the most intense^17^, for three consecutive winters (2017, 2018 and 2019). We implemented the experiment on 18 individuals, including five recaptures and two deployments spanning two winters, leading to a total of 25 animal-years (21 adults: 15 females, 6 males; 4 yearlings: 2 females, 2 males; sample size *n*=4, 11 and 10 in 2017, 2018 and 2019 respectively; see Supplementary S2 for details). The animal-year was our sampling unit, on the assumption that the same individual may respond independently to manipulations in different years. Roe deer were captured using baited box traps (n=16) or net drives (n=3), and were fitted with GPS-GSM radio collars programmed to acquire hourly GPS locations for a year, after which they were released via a drop-off mechanism. Captures and marking were performed complying with ethical and welfare rules, under authorization of the Wildlife Committee of the Autonomous Province of Trento (Resolution of the Provincial Government n. 602, under approval of the Wildlife Committee of 20/09/2011, and successive integration approved on the 23/04/2015). Radio-collared roe deer moved an average of 61.2 m per hour. This value of the average hourly movement distance *(l)* was subsequently utilized in the analyses described below.

For all captured animals, we assumed a post-capture response in ranging behaviour. We therefore considered the first re-visitation of the capture location as the sign of resettlement in the original range and we used this time as onset of the experimental pre-closure phase. Although not all the individuals were manipulated at the same time, we avoided interference between capture operations and FS manipulations, and between co-occurring different manipulation phases (i.e., ensuring that co-occurring manipulations occurred in separate areas).

During the *pre-closure phase*, we assessed the use of FS by radio-collared roe deer. We identified the “manipulated” FS (M) for each individual as the site with the largest number of locations within a radius *l* during this initial phase, and considered it as the FS to which an individual is most familiar. During the *closure phase*, corn was made inaccessible at M for a duration of about 15 days, depending on personnel availabilities (min=14.0 days, max=18.1, mean=15.5). M was then re-opened, thereby initiating the *post-closure phase*. During both pre- and post-closure phases, corn was available *ad libitum* at M. All “alternate” (A) managed FS – i.e., that were provisioned at least once in the month prior to the experiment – had corn available *ad libitum* throughout the duration of the experiment.

### Data preparation

To ensure meaningful comparisons between animal-years, we homogenized the durations of each experimental phase to the minimum length of the closure phase in our sample (i.e., 14 days). Specifically, we truncated the movement data by removing initial excess positions for the pre-closure and closure phases, and terminal excess positions for the post-closure phase. GPS acquisition success was extremely high (99.57 % during the experiment) and we did not interpolate missing fixes in the collected data.

The analyses of space-use and movement behaviour were based on spatially-explicit, raw movement trajectories. The analyses of resource use, instead, relied on spatially-implicit, state time series derived from the underlying movement data. To this end, we created an initial time series, for each animal-year, by intersecting the relocations with three spatial domains: vegetation (the matrix; V), manipulated FS (M) and alternate FS (A). We converted FS locations (M and A) into areas by buffering them. To investigate the sensitivity of buffer choice we considered six buffer sizes: *l* (i.e., 61.2 m) multiplied by 0.5, 1, 1.5, 2, 3 and 4. We associated all locations falling outside M and A to the state V. The three-state time series was then converted into three single-state presence/absence time series.

### Preference for feeding sites

We calculated each individual’s preference for FS (*h*_*FS*_) as the relative use of FS over natural vegetation during the pre-closure phase (i.e., the proportion of GPS fixes classified as either M or A). Because preference is considered to be temporally dynamic^30^, we chose to evaluate *h*_*FS*_ for each year separately in case individuals were manipulated in two separate winters. This reasoning allowed to account for the influence of individual condition and of the relative quality and quantity of vegetation resources on *h*_*FS*_. We included *h*_*FS*_ in all space-use, movement, and resource use analyses described below.

The variability of *h*_*FS*_ across animal-years was maximal when FS attendance was defined as a GPS location within a distance equal to the population mean hourly step length *l* i.e., 61.2 m from the FS (interquartile range=0.278, mean=0.343; Supplementary S3: Table S1). Accordingly, the results described below are based on this definition (see Supplementary S7 for a sensitivity analysis). At this scale, *h*_*FS*_ did not differ consistently between sex (mean for females=0.346; mean for males=0.336; t-test: p-value=0.901).

### Analysis

We analysed how the experimental manipulation, and its interaction with both preference for FS and sex, affected roe deer space-use, movement behaviour, and resource use.

#### Space-use

We assessed the changes of home range and core area sizes (P1.1), and of space-use overlap (P1.2, P3.1) between experimental phases. We calculated utilization distributions (UD)^31^ for each animal-year and experimental phase using a Gaussian kernel density estimation. After visual inspection, we chose to compute the UDs at a spatial resolution of 10 m and with a fixed bandwidth set to half the average hourly movement distance (i.e., *l*/2=30.6 m).

For home range and core area sizes, we calculated the area (in hectares) corresponding to the 95% and 50% UD contours, respectively, during each experimental phase (*Phase*; three levels; reference level: *Pre-closure*). We then analysed the log-transformed areas using a linear mixed-effect model (LMM) with five fixed effects: *Phase*, *h*_*FS*_, *Sex* (categorical predictor; reference level: *Female*), and two interaction terms (*Phase*:*h*_*FS*_ and *Phase:Sex*). We included animal-year (*ind*) as random effect (intercept). In all analyses, interaction terms were dropped when statistically non-significant (p-value>0.05).

For space-use patterns, we estimated the overlaps for three pairs of UDs – pre- and post-closure, pre-closure and closure, and closure and post-closure (*Contrast*; three levels; reference level: Pre-/Closure) – using the volume of intersection statistic (VI)^32^. VI ranges from 0 (no overlap) to 1 (complete overlap). We analysed the logit-transformed overlaps using an LMM with *Contrast*, *h*_*FS*_, *Sex*, *Contrast*:*h*_*FS*_ and *Contrast:Sex* as fixed effects, and *ind* as random intercept.

#### Movement behaviour

We investigated the movement responses of roe deer to the experiment (P1.3) by analysing the changes in hourly step length (Euclidean distance between two successive relocations) and turning angle *θ*_*t*_ (angle between two successive movement steps). Turning angles range between −π and π, and were symmetric around 0. We analysed the log-transformed step length, *s*_*t*_, and the logit-transformed absolute turning angle, 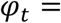 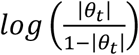 using LMMs with *Phase*, *h*_*FS*_, *Sex*, *Phase*:*h*_*FS*_ and *Phase:Sex* as fixed effects, and *ind* as random intercept. Because step length was characterized by strong serial autocorrelation at short temporal lags and at circadian periodicities (a common pattern in animal movement trajectories^33^), we also included step length measured at lags 1, 2 and 24 h (i.e., *s*_*t*−1_, *s*_*t*−2_, *s*_*t*−24_) as fixed effects to reduce the autocorrelation of the model residuals.

#### Resource use

To test whether the experiment led to a transitory change in resource use (P1.4a-b, P3.2), we fitted separate mixed-effect logistic regression models to the three single-state presence/absence time series (*u*_*M*,*t*_, *u*_*A*,*t*_ and *u*_*V*,*t*_) using *Phase*, *h*_*FS*_, *Sex*, *Phase*:*h*_*FS*_ and *Phase:Sex* as fixed effects, and *ind* as random intercept. The pre-closure level for *Phase* was dropped for *u*_*V*_ to avoid circularity (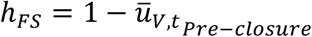). We also included the response variables measured at lags 1, 2 and 24 h (e.g., *u*_*M*,*t*−1_, *u*_*M*,*t*−2_, *u*_*M*,*t*−24_) as fixed effects to reduce the autocorrelation of the model residuals. However, for the sake of conciseness and clarity, we omit these response lags when visualizing resource use predictions. Because the model results were consistent regardless of the inclusion of the response lags (Supplementary S6: Tables S1, S3), this decision had no impact on the interpretation. Two animal-years were excluded from the analyses of resource use due to the absence of suitable A-state: F4-2017 did not seem to have visited any alternate FS (A) prior to the experiment; and F16-2016 had two distinct, highly-used FS during pre-closure, but only the second most visited FS could be manipulated (due to stakeholder acceptance). While the use of A was more variable when including these two outliers, the general patterns remained unchanged (Supplementary S6: Tables S1, S4).

#### Software

All analyses were conducted in the R environment^34^. We used the packages adehabitatLT and adehabitatHR^35^ for the spatial analyses, fitted all mixed-effect models via Maximum Likelihood with the package lme4^36^, and obtained the coefficients of determination using MuMin^37^.

## Acknowledgements

N. Ranc was supported by Harvard University Graduate Fellowship and Fondazione Edmund Mach International Doctoral Programme Fellowship. F. Cagnacci was supported by the Sarah and Daniel Hrdy Fellowship 2015-2016 at Harvard University OEB during part of the development of this manuscript. We are grateful to all the people involved in the roe deer captures and data collection, and especially M.B. Almeida, P. Bonanni, A. Corradini, J. De Groeve, G. Lilli, S. Mumme, S. Nicoloso, D. Righetti, M. Rocca, M. Salvatori, M. Sanchez and P. Semenzato. We thank J.W. Cain, B. Ölveczky, N. Pierce, all the members of the Cagnacci lab and of the Moorcroft lab, and the graduate students of the Harvard Statistics Consulting Service for their valuable suggestions. We also thank the Wildlife and Forestry Service of the Autonomous Province of Trento and the Trentino Hunting Association (ACT) for their support. Finally, we express our gratitude to the hunters of Albiano, Civezzano, Fornace and Trento Nord without whose help this study could not be conducted.

