## Supplementary Materials for "Preference and familiarity mediate spatial responses of a large herbivore to experimental manipulation of resource availability"

**Supplementary S1: Study area description**

The study area is located in the north-eastern Italian Alps (Argentario range, in Val di Cembra and Valsugana; Autonomous Province of Trento). It covers 44.91 $km^2$ and ranges between 300 and 1,000 m a.s.l. The topography is generally mild, but steeper slopes (> 30°) occur in the northern portion. The climate is continental and characterized by a mean temperature of 1.0 °C in January and 21.0 °C in July, and a mean annual rainfall of 966 mm (average 2000-2018; http://www.meteotrentino.it). There is occasional snow cover between December and March.

The study area is covered by 80.0 % forest (High-Resolution Raster Later Tree Cover Density; EEA 2012; see eea.europa.eu), mostly as secondary growth stands interspersed with small pastures. The forests are dominated by *Pinus sylvestris* with abundant shrub undergrowth, and by mixed stands of *Fagus sylvatica*, *Picea abies* and *Abies alba* and, to a lower extent, by *Quercus petraea* stands. It is a multi-use landscape that is managed for selective logging forestry, localized mining, small-scale agriculture, and recreational activities. Two paved roads and a dense network of dirt roads and trails connect small towns located at lower elevation. Two ungulates are present in the area: roe deer, which is most prevalent, and red deer, whose presence is more sporadic but is increasing (7-8 ind. $km^{-2}$ for roe deer and <1 ind. $km^{-2}$ for red deer; ref. values from Autonomous Province of Trento Wildlife Office). The largest natural predator of roe deer (especially fawns) in this landscape is the red fox (*Vulpes vulpes*).

Supplemental forage is provided year-round at designated FS (official authorization: "Autonomous Province of Trento order n. 2852/2013") by private hunters, with support from the Trentino Hunting Association. FS are managed for roe deer but are also attended sporadically by red deer (*Cervus elaphus*), European badgers (*Meles meles*), red squirrels (*Sciurus vulgaris*), rodents (*Apodemus sp*., *Microtus sp*.), European jays (*Garrulus glandarius*) and wood pigeons (*Columba palumbus*)

**Supplementary S2: Animal captures and tracking**

Between November 2016 and February 2019, we captured and marked 37 roe deer using wooden box traps baited with corn near feeding sites in winter (n = 33) and net drives in spring and fall (n = 5). Of these captured individuals, 26 (yearlings and adults, or fawns captured after March) were fitted with GPS-GSM radio collars (VECTRONIC Aerospace GmbH; models GPS Plus, Vertex Plus or Vertex Lite). Nine individuals were recaptured in two separate years (n=7) or had data spanning two subsequent winters (n=2), thereby leading to a total of 35 animal-years (28 adults: 21 females, 7 males; 7 yearlings/fawns: 5 females, 2 male). Two collar batteries failed prior to this period. In addition, prerequisites for performing the experimental manipulation on an animal-year were: (i) spatial overlap between the animal-year movement trajectory and a FS, defined here as at least 10 relocations within a radius $l$ (mean hourly step length i.e., 61.2 m) of any managed FS, during a two-week period (i.e., the pre-closure) and (ii) possibility to alter the FS management, after explicit agreement with its private owner, which led to the exclusion of eight animal-years from the experiment. In light of the above considerations, **we retained 25 animal-years** (21 adults: 15 females, 6 males; 4 yearlings: 2 females, 2 males; n=4 in 2017, n=11 in 2018 and n=10 in 2019) for the experimental manipulation. One animal died (F4-2018), and another had a prolonged series of missing fixes (F28-2019) during the third phase of the experimental manipulation, so we excluded two post-closure phases.

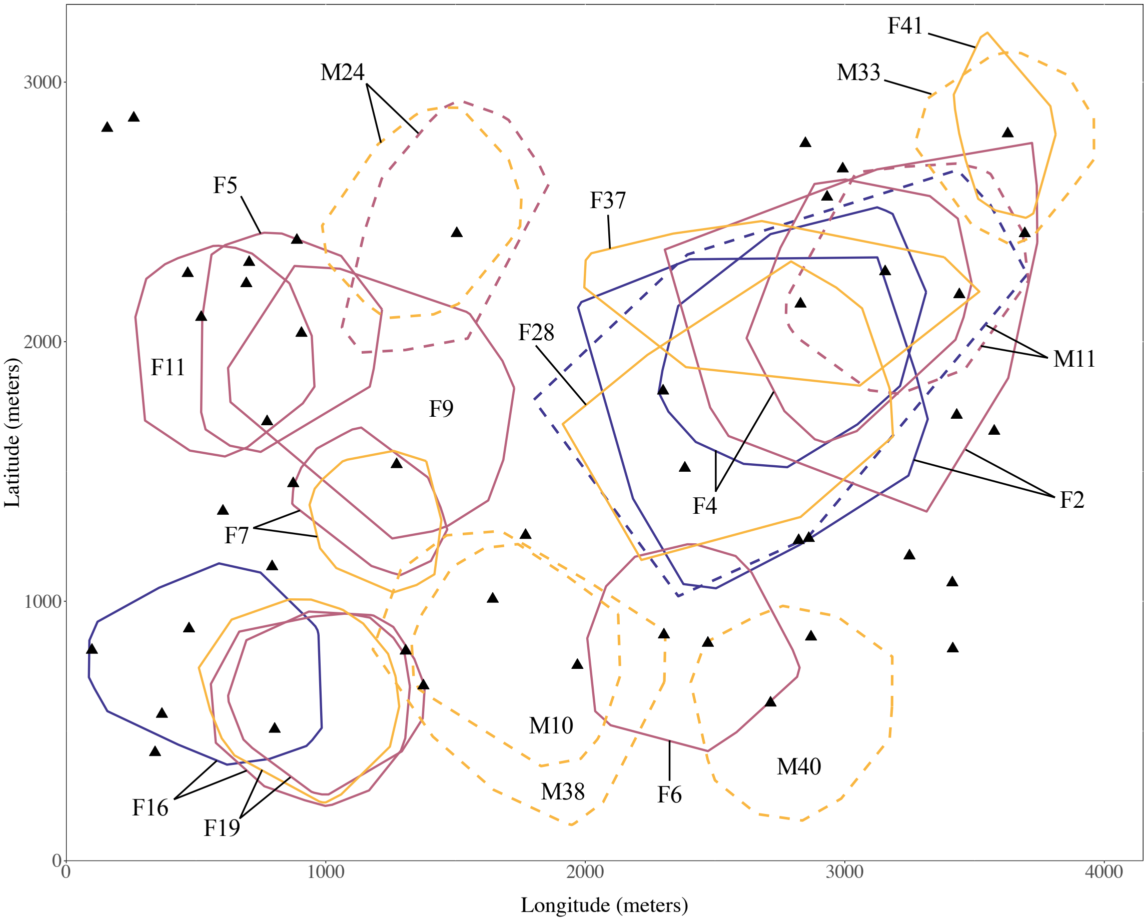


Figure S1. Spatial distribution of the roe deer included in the experiment. The area occupied by each twenty-five animal-years is plotted as a 95 % minimum convex polygon (2017: blue; 2018: burgundy; 2019: orange). Female home ranges are displayed as solid lines and males as dashed lines. All managed feeding sites (FS i.e., either M or A) are identified as black triangles.

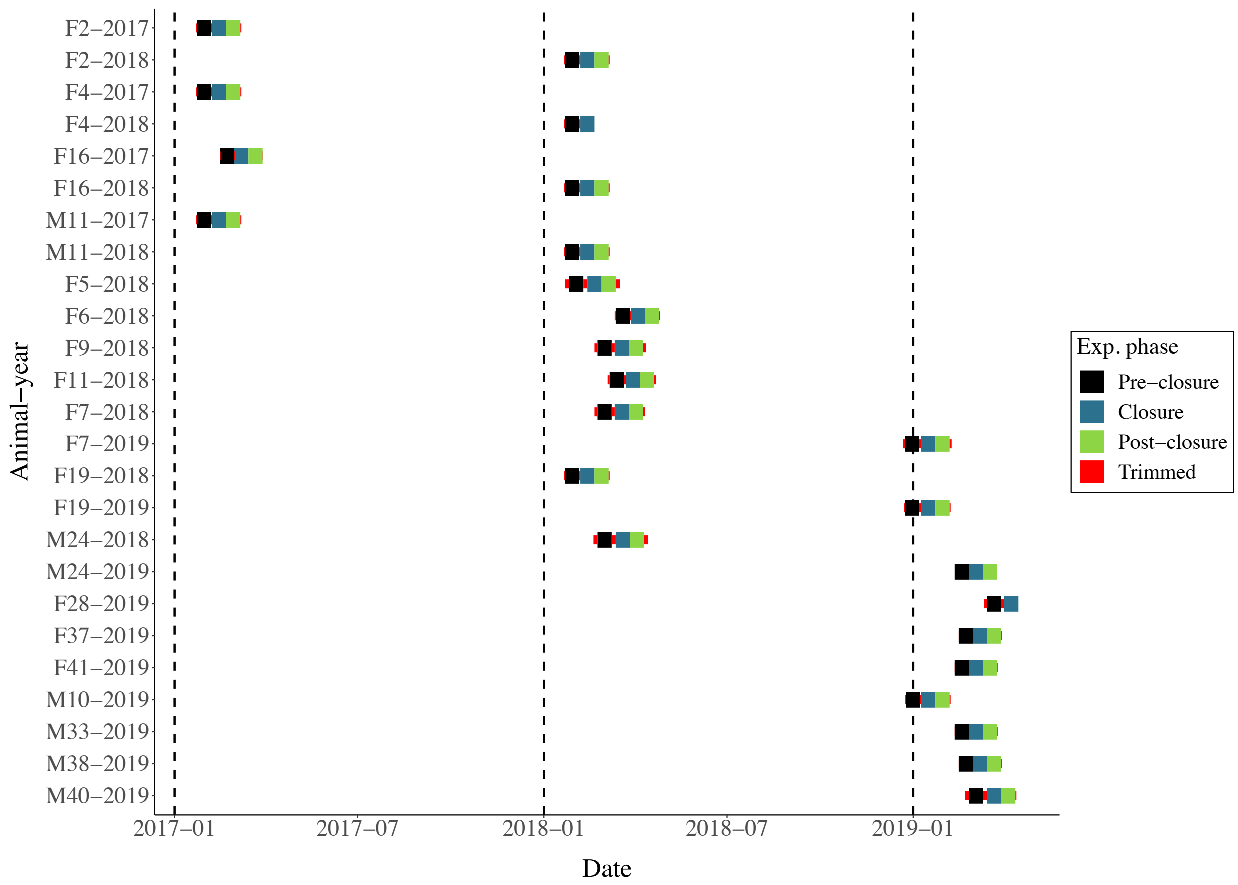


Figure S2. Monitoring history of the roe deer included in the experiment. To ensure comparability among animal-years, the initial excess positions for the pre-closure and closure phases and terminal excess positions for post-closure phase were trimmed. The post-closure phases of F4-2018 and F28-2019 have been excluded from the analyses due to a mortality case and a high proportion of missing fixes, respectively.

**Supplementary S3: Individual variability in feeding site preference**

Table S1. Preference for feeding sites ($h_{FS}$) calculated for each animal-year (ID-Year) and based on six feeding site buffer sizes: mean step length of roe deer, $l$, multiplied by 0.5, 1, 1.5, 2, 3 and 4 (i.e., 30.6, 61.2, 91.8, 122.4, 183.6 and 244.8 m, respectively). The inter-individual variability in $h_{FS}$ (interquartile range and standard deviation) is maximum for a buffer of $l$.

| | Buffer size – multiple of the mean step length ($l$) | | | | | |
|---|---|---|---|---|---|---|
| Animal-year ID | 0.5 | 1.0 | 1.5 | 2.0 | 3.0 | 4.0 |
| F2-2017 | 0.352 | **0.752** | **0.797** | **0.830** | **0.904** | **0.931** |
| F2-2018 | 0.215 | 0.540 | 0.585 | 0.618 | 0.749 | 0.866 |
| F4-2017 | 0.155 | 0.615 | 0.693 | 0.749 | 0.863 | 0.961 |
| F4-2018 | 0.084 | 0.499 | 0.597 | 0.633 | 0.693 | 0.848 |
| F16-2017 | 0.301 | 0.418 | 0.537 | 0.624 | 0.809 | 0.952 |
| F16-2018 | 0.149 | 0.281 | 0.325 | 0.472 | 0.710 | 0.863 |
| M11-2017 | 0.170 | 0.639 | 0.734 | 0.755 | 0.845 | 0.940 |
| M11-2018 | 0.164 | 0.493 | 0.534 | 0.603 | 0.699 | 0.779 |
| F5-2018 | 0.101 | 0.242 | 0.310 | 0.478 | 0.710 | 0.872 |
| F6-2018 | 0.149 | 0.242 | 0.331 | 0.379 | 0.481 | 0.630 |
| F9-2018 | 0.069 | 0.110 | **0.158** | **0.218** | 0.382 | 0.633 |
| F11-2018 | 0.078 | 0.290 | 0.355 | 0.394 | 0.481 | 0.591 |
| F7-2018 | 0.084 | 0.233 | 0.388 | 0.469 | 0.618 | 0.761 |
| F7-2019 | 0.051 | 0.152 | 0.287 | 0.376 | 0.531 | 0.782 |
| F19-2018 | 0.140 | 0.296 | 0.346 | 0.451 | 0.672 | 0.848 |
| F19-2019 | 0.107 | 0.140 | 0.188 | 0.218 | 0.388 | 0.639 |

| | | | | | | |
|---|---|---|---|---|---|---|
| M24-2018 | 0.137 | 0.412 | 0.555 | 0.687 | 0.803 | 0.899 |
| M24-2019 | 0.063 | 0.215 | 0.424 | 0.543 | 0.696 | 0.761 |
| F28-2019 | **0.358** | 0.570 | 0.630 | 0.710 | 0.791 | 0.881 |
| F37-2019 | **0.036** | 0.084 | 0.170 | 0.496 | 0.737 | 0.896 |
| F41-2019 | 0.140 | 0.421 | 0.484 | 0.528 | 0.869 | 0.943 |
| M10-2019 | 0.218 | 0.296 | 0.373 | 0.451 | 0.567 | 0.693 |
| M33-2019 | 0.200 | 0.394 | 0.454 | 0.549 | 0.773 | 0.890 |
| M38-2019 | 0.036 | **0.075** | 0.188 | 0.334 | 0.448 | 0.606 |
| M40-2019 | 0.104 | 0.164 | 0.224 | 0.296 | **0.379** | **0.558** |
| Interquartile range | 0.087 | **0.278** | 0.245 | 0.230 | 0.260 | 0.203 |
| Mean | 0.147 | 0.343 | 0.427 | 0.514 | 0.664 | 0.801 |
| Standard deviation | 0.089 | **0.188** | 0.183 | 0.164 | 0.163 | 0.128 |
| Minimum | 0.036 | 0.075 | 0.158 | 0.218 | 0.379 | 0.558 |
| Maximum | 0.358 | 0.752 | 0.797 | 0.830 | 0.904 | 0.961 |

### Supplementary S4: Supplementary results – space-use models

The final models were obtained by removing non-significant interaction terms from the full models.

*Home range and core area sizes*

Table S1. Summary of the final model for home range size (95% UD). The model includes experimental phase (*Phase*; reference level: *Pre-closure*), preference for feeding sites ($h_{FS}$) and their interaction as fixed effects, and animal-year as a random intercept.

| | Estimate | Std. Error | df | t value | p-value |
|---|---|---|---|---|---|
| (Intercept) | 3.392 | 0.162 | 45.118 | 21.002 | <0.001*** |
| *PhaseClosure* | -0.001 | 0.150 | 48.269 | -0.005 | 0.996 |
| *PhasePost-closure* | -0.138 | 0.151 | 48.395 | -0.914 | 0.365 |
| $h_{FS}$ | -0.532 | 0.415 | 45.118 | -1.282 | 0.206 |
| *PhaseClosure*:$h_{FS}$ | 0.845 | 0.385 | 48.269 | 2.195 | 0.033* |
| *PhasePost-closure:*$h_{FS}$ | 0.879 | 0.399 | 48.870 | 2.205 | 0.032* |

| | Std. Dev | | $R^2$ |
|---|---|---|---|
| Random effect | 0.289 | Marginal | 0.122 |
| Residual | 0.251 | Conditional | 0.622 |

Table S2. Summary of the final model for home range size (50% UD). The model includes experimental phase (*Phase*; reference level: *Pre-closure*), preference for feeding sites ($h_{FS}$) and their interaction as fixed effects, and animal-year as a random intercept.

| | Estimate | Std. Error | df | t value | p-value |
|---|---|---|---|---|---|
| (Intercept) | 1.930 | 0.181 | 65.921 | 10.684 | <0.001*** |
| *PhaseClosure* | -0.106 | 0.223 | 48.722 | -0.478 | 0.635 |
| *PhasePost-closure* | -0.101 | 0.224 | 48.970 | -0.450 | 0.655 |
| $h_{FS}$ | -2.096 | 0.464 | 65.921 | -4.514 | <0.001*** |
| *PhaseClosure*:$h_{FS}$ | 1.572 | 0.572 | 48.722 | 2.747 | 0.008** |
| *PhasePost-closure:*$h_{FS}$ | 1.179 | 0.591 | 49.914 | 1.997 | 0.051(*) |

| | Std. Dev |
|---|---|
| Random effect | 0.209 |
| Residual | 0.373 |

| | $R^2$ |
|---|---|
| Marginal | 0.351 |
| Conditional | 0.506 |

Table S3. Summary of the full model for home range size (95% UD). The model includes experimental phase (*Phase*; reference level: *Pre-closure*), preference for feeding sites ($h_{FS}$), *Sex* (reference level: female, *F*), and the interactions of *Phase* with both $h_{FS}$ and *Sex* as fixed effects, and animal-year as a random intercept.

| | Estimate | Std. Error | df | t value | p-value |
|---|---|---|---|---|---|
| (Intercept) | 3.344 | 0.163 | 46.998 | 20.470 | <0.001*** |
| *PhaseClosure* | -0.043 | 0.157 | 48.197 | -0.275 | 0.785 |
| *PhasePost-closure* | -0.184 | 0.157 | 48.238 | -1.167 | 0.249 |
| $h_{FS}$ | -0.523 | 0.397 | 46.998 | -1.318 | 0.194 |
| *Sex* | 0.141 | 0.157 | 46.998 | 0.903 | 0.371 |

| | | | | | |
|---|---|---|---|---|---|
| *PhaseClosure*:$h_{FS}$ | 0.853 | 0.381 | 48.197 | 2.237 | 0.030* |
| *PhasePost-closure:*$h_{FS}$ | 0.863 | 0.395 | 48.858 | 2.186 | 0.034* |
| *PhaseClosure*:*SexM* | 0.124 | 0.151 | 48.197 | 0.822 | 0.415 |
| *PhasePost-closure*:*SexM* | 0.146 | 0.153 | 48.473 | 0.954 | 0.345 |
| | Std. Dev | | | $R^2$ | |
| Random effect | 0.268 | | Marginal | 0.200 | |
| Residual | 0.248 | | Conditional | 0.630 | |

Table S4. Summary of the full model for core area size (50% UD). The model includes experimental phase (*Phase*; reference level: *Pre-closure*), preference for feeding sites ($h_{FS}$), *Sex* (reference level: female, *F*), and the interactions of *Phase* with both $h_{FS}$ and *Sex* as fixed effects, and animal-year as a random intercept.

| | Estimate | Std. Error | df | t value | p-value |
|---|---|---|---|---|---|
| (Intercept) | 1.925 | 0.182 | 67.324 | 10.563 | <0.001*** |
| *PhaseClosure* | -0.235 | 0.229 | 48.720 | -1.027 | 0.309 |
| *PhasePost-closure* | -0.185 | 0.229 | 48.798 | -0.808 | 0.423 |
| $h_{FS}$ | -2.095 | 0.443 | 67.324 | -4.734 | <0.001*** |
| *Sex* | 0.015 | 0.175 | 67.324 | 0.084 | 0.933 |
| *PhaseClosure*:$h_{FS}$ | 1.597 | 0.555 | 48.720 | 2.876 | 0.006** |
| *PhasePost-closure:*$h_{FS}$ | 1.162 | 0.573 | 49.976 | 2.028 | 0.048* |
| *PhaseClosure*:*SexM* | 0.375 | 0.219 | 48.720 | 1.710 | 0.094(*) |
| *PhasePost-closure*:*SexM* | 0.263 | 0.222 | 49.242 | 1.185 | 0.242 |

| | Std. Dev | | $R^2$ | |
|---|---|---|---|---|
| Random effect | 0.188 | | Marginal | 0.411 |
| Residual | 0.362 | | Conditional | 0.536 |

*Space-use overlap*

Table S5. Summary of the final model for space-use overlap. The model includes experimental contrast (*Contrast*; reference level: *Pre-c./Closure*), preference for feeding sites ($h_{FS}$), and the interaction of *Contrast* with $h_{FS}$ as fixed effects, and animal-year as a random intercept.

| | Estimate | Std. Error | df | t value | p-value |
|---|---|---|---|---|---|
| (Intercept) | 0.096 | 0.225 | 66.109 | 0.428 | 0.670 |
| *ContrastClosure/Post-c.* | 0.119 | 0.286 | 47.785 | 0.414 | 0.680 |
| *ContrastPre-c./Post-c.* | -0.045 | 0.286 | 47.785 | -0.156 | 0.877 |
| $h_{FS}$ | -2.073 | 0.578 | 66.109 | -3.590 | <0.001** |
| *ContrastClosure/Post-c.*:$h_{FS}$ | 1.179 | 0.753 | 49.192 | 1.565 | 0.124 |
| *ContrastPre-c./Post-c.*:$h_{FS}$ | 2.366 | 0.753 | 49.192 | 3.141 | 0.003** |

| | Std. Dev | | $R^2$ | |
|---|---|---|---|---|
| Random effect | 0.238 | | Marginal | 0.372 |
| Residual | 0.476 | | Conditional | 0.498 |

Table S6. Summary of the full model for space-use overlap. The model includes experimental contrast (*Contrast*; reference level: *Pre-c./Closure*), preference for feeding sites ($h_{FS}$), *Sex* (reference level: female, *F*), and the interactions of *Contrast* with both $h_{FS}$ and *Sex* as fixed effects, and animal-year as a random intercept.

| | Estimate | Std. Error | df | t value | p-value |
|---|---|---|---|---|---|
| (Intercept) | 0.061 | 0.233 | 64.911 | 0.261 | 0.795 |
| *ContrastClosure/Post-c.* | 0.085 | 0.291 | 47.529 | 0.293 | 0.771 |
| *ContrastPre-c./Post-c.* | 0.082 | 0.291 | 47.529 | 0.283 | 0.778 |
| $h_{FS}$ | -2.066 | 0.567 | 64.911 | -3.646 | <0.001** |
| *Sex* | 0.103 | 0.224 | 64.911 | 0.461 | 0.647 |
| *ContrastClosure/Post-c.*:$h_{FS}$ | 1.152 | 0.728 | 49.210 | 1.583 | 0.120 |
| *ContrastPre-c./Post-c.*:$h_{FS}$ | 2.390 | 0.728 | 49.210 | 3.283 | 0.002** |
| *ContrastClosure/Post-c.:SexM* | 0.114 | 0.282 | 48.162 | 0.403 | 0.689 |
| *ContrastPre-c./Post-c.:SexM* | -0.395 | 0.282 | 48.162 | -1.401 | 0.168 |

| | Std. Dev |
|---|---|
| Random effect | 0.248 |
| Residual | 0.459 |

| | $R^2$ |
|---|---|
| Marginal | 0.396 |
| Conditional | 0.533 |

### Supplementary S5: Supplementary results – movement models

The final models were obtained by removing non-significant interaction terms from the full models.

*Step length*

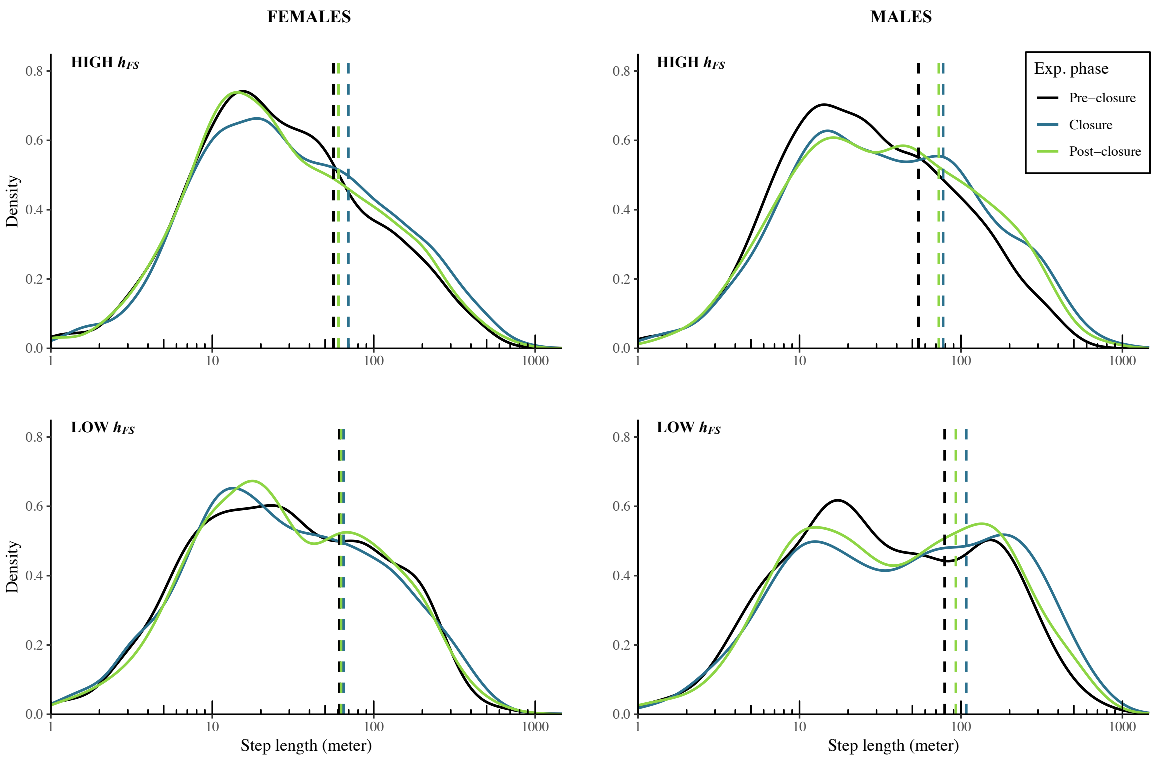


Figure S1. Changes in step length distribution across the three experimental phases (colour) for females (left panels) and males (right panels) with high preference for feeding sites (larger or equal to the sample median i.e., $h_{FS} \geq 0.29$; top panels) and low $h_{FS}$ ($h_{FS} < 0.29$; bottom panels). Vertical dashed lines indicate step length means.

Table S1. Summary of the final model for step length ($s_t$). The model includes experimental phase (*Phase*; reference level: *Pre-closure*), preference for feeding sites ($h_{FS}$), *Sex* (reference level: female, *F*), the interactions of *Phase* with both $h_{FS}$ and *Sex*, and the step length at lags 1, 2 and 24 hours ($s_{t-1}$, $s_{t-2}$ and $s_{t-24}$) as fixed effects, and animal-year as a random intercept.

| | Estimate | Std. Error | df | t value | p-value |
|---|---|---|---|---|---|
| (Intercept) | 2.537 | 0.059 | 90.598 | 42.764 | <0.001*** |
| *PhaseClosure* | -0.025 | 0.043 | 23950.039 | -0.568 | 0.570 |
| *PhasePost-closure* | 0.007 | 0.043 | 23962.736 | 0.172 | 0.864 |
| $h_{FS}$ | -0.319 | 0.120 | 43.862 | -2.659 | 0.011* |
| *Sex* | 0.052 | 0.047 | 43.964 | 1.089 | 0.282 |
| *PhaseClosure*:$h_{FS}$ | 0.244 | 0.105 | 23950.469 | 2.328 | 0.020* |
| *PhasePost-closure*:$h_{FS}$ | 0.041 | 0.108 | 23510.633 | 0.379 | 0.704 |
| *PhaseClosure*:*SexM* | 0.128 | 0.042 | 23951.008 | 3.084 | 0.002** |
| *PhasePost-closure:SexM* | 0.125 | 0.042 | 23935.032 | 2.957 | 0.003** |
| $s_{t-1}$ | 0.284 | 0.006 | 23972.255 | 44.688 | <0.001*** |
| $s_{t-2}$ | -0.148 | 0.006 | 23973.995 | -23.405 | <0.001*** |
| $s_{t-24}$ | 0.128 | 0.006 | 23969.001 | 20.892 | <0.001*** |

| | Std. Dev |
|---|---|
| Random effect | 0.088 |
| Residual | 1.991 |

| | $R^2$ |
|---|---|
| Marginal | <0.01 |
| Conditional | <0.01 |

*Turn angle*

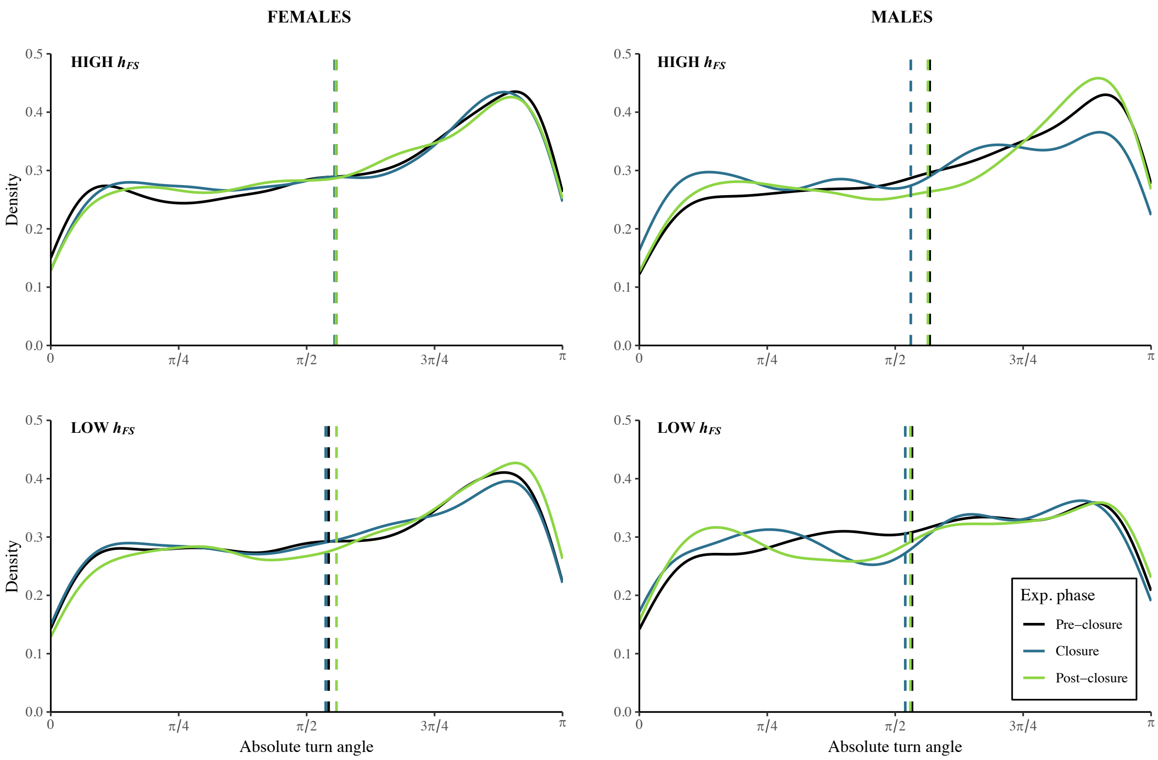


Figure S2. Changes in absolute turn angle distribution across the three experimental phases (colour) for females (left panels) and males (right panels) with high preference for feeding sites (larger or equal to the sample median i.e., $h_{FS} \geq 0.29$; top panels) and low $h_{FS}$ ($h_{FS} < 0.29$; bottom panels). Vertical dashed lines indicate absolute turn angle means.

Table S2. Summary of the final model for the absolute turn angle ($\varphi_t$). The model includes experimental phase (*Phase*; reference level: *Pre-closure*), preference for feeding sites ($h_{FS}$), *Sex* (reference level: female, *F*) and the interaction of *Phase* with *Sex* as fixed effects, and animal-year as a random intercept

| | Estimate | Std. Error | df | t value | p-value |
|---|---|---|---|---|---|
| (Intercept) | 0.194 | 0.054 | 35.811 | 3.623 | <0.001*** |
| *PhaseClosure* | -0.033 | 0.038 | 23771.982 | -0.869 | 0.385 |
| *PhasePost-closure* | 0.067 | 0.039 | 21733.369 | 1.714 | 0.087(*) |
| $h_{FS}$ | 0.371 | 0.119 | 24.934 | 3.118 | 0.005** |
| *Sex* | 0.059 | 0.060 | 68.683 | 0.984 | 0.329 |
| *PhaseClosure*:*SexM* | -0.187 | 0.067 | 23772.703 | -2.783 | 0.005** |
| *PhasePost-closure:SexM* | -0.113 | 0.068 | 23619.032 | -1.662 | 0.096(*) |

| | Std. Dev | | $R^2$ |
|---|---|---|---|
| Random effect | 0.088 | Marginal | <0.01 |
| Residual | 1.992 | Conditional | <0.01 |

Table S3. Summary of the full model for the absolute turn angle ($\varphi_t$). The model includes experimental phase (*Phase*; reference level: *Pre-closure*), preference for feeding sites ($h_{FS}$), *Sex* (reference level: female, *F*), and the interactions of *Phase* with both $h_{FS}$ and *Sex* as fixed effects, and animal-year as a random intercept.

| | Estimate | Std. Error | df | t value | p-value |
|---|---|---|---|---|---|
| (Intercept) | 0.132 | 0.063 | 69.488 | 2.089 | 0.040* |
| *PhaseClosure* | 0.063 | 0.070 | 23771.135 | 0.904 | 0.366 |
| *PhasePost-closure* | 0.161 | 0.070 | 23788.293 | 2.308 | 0.021* |
| $h_{FS}$ | 0.551 | 0.152 | 68.487 | 3.614 | <0.001*** |

| | | | | | |
|---|---|---|---|---|---|
| *Sex* | 0.062 | 0.060 | 69.158 | 1.023 | 0.310 |
| *PhaseClosure*:$h_{FS}$ | -0.276 | 0.168 | 23770.632 | -1.638 | 0.101 |
| *PhasePost-closure*:$h_{FS}$ | -0.278 | 0.174 | 22276.612 | -1.596 | 0.110 |
| *PhaseClosure*:*SexM* | -0.190 | 0.067 | 23772.720 | -2.829 | 0.005** |
| *PhasePost-closure:SexM* | -0.113 | 0.068 | 23579.013 | -1.667 | 0.096(*) |

| | Std. Dev | | $R^2$ |
|---|---|---|---|
| Random effect | 0.088 | Marginal | <0.01 |
| Residual | 1.991 | Conditional | <0.01 |

**Supplementary S6: Supplementary results – resource use models**

The final models were obtained by removing non-significant interaction terms from the full models.

Table S1. Summary of the final models for the use of the manipulated feeding site ($u_{M,t}$), alternate feeding sites ($u_{A,t}$) and vegetation ($u_{V,t}$). The models include experimental phase (*Phase*; reference level: *Pre-closure*), preference for feeding sites ($h_{FS}$), *Sex* (reference level: female, *F;* only retained for $u_{A,t}$), the interactions of *Phase* with both $h_{FS}$ and *Sex* (only retained for $u_{A,t}$), and the resource variables at lags 1, 2 and 24 hours (e.g., $u_{M,t-1}$, $u_{M,t-2}$ and $u_{M,t-24}$) as fixed effects, and animal-year as a random intercept. For the vegetation model, the data included only the Closure and Post-closure phases since the average $u_{V,t}$ during pre-closure was used to calculate $h_{FS}$. The reference levels used for *Phase* were *Pre-closure* for $u_{M,t}$, and *Closure* for $u_{V,t}$.

| | Manipulated feeding site (M) | | | |
|---|---|---|---|---|
| | Estimate | Std. Error | z value | p-value |
| (Intercept) | -3.325 | 0.112 | -29.804 | <0.001*** |
| *PhaseClosure* | -0.645 | 0.172 | -3.740 | <0.001*** |
| *PhasePost-closure* | -0.102 | 0.126 | -0.809 | 0.418 |
| $h_{FS}$ | 1.723 | 0.288 | 5.990 | <0.001*** |
| *PhaseClosure*:$h_{FS}$ | -1.657 | 0.459 | -3.606 | <0.001*** |
| *PhasePost-closure:*$h_{FS}$ | -0.473 | 0.325 | -1.456 | 0.145 |
| $u_{M,t-1}$ | 3.155 | 0.065 | 48.555 | <0.001*** |

| | | | | |
|---|---|---|---|---|
| $u_{M,t-2}$ | 0.900 | 0.068 | 13.286 | <0.001*** |
| $u_{M,t-24}$ | 0.739 | 0.062 | 11.917 | <0.001*** |
| | Std. Dev | | | $R^2$ |
| Random effect | 0.161 | | Marginal | 0.346 |
| Residual | 1.000 | | Conditional | 0.349 |

| | Alternate feeding sites (A) | | | |
|---|---|---|---|---|
| | Estimate | Std. Error | z value | p-value |
| (Intercept) | -6.032 | 0.379 | -15.913 | <0.001*** |
| *PhaseClosure* | 2.191 | 0.297 | 7.38 | <0.001*** |
| *PhasePost-closure* | 1.663 | 0.308 | 5.395 | <0.001*** |
| $h_{FS}$ | 3.869 | 0.829 | 4.668 | <0.001*** |
| *Sex* | -0.917 | 0.372 | -2.464 | 0.014* |
| *PhaseClosure*:$h_{FS}$ | -1.726 | 0.571 | -3.022 | 0.003** |
| *PhasePost-closure:*$h_{FS}$ | -1.831 | 0.594 | -3.081 | 0.002** |
| *PhaseClosure:SexM* | 0.529 | 0.292 | 1.815 | 0.069(*) |
| *PhasePost-closure:SexM* | 0.855 | 0.302 | 2.835 | 0.005** |
| $u_{A,t-1}$ | 2.993 | 0.081 | 36.935 | <0.001*** |
| $u_{A,t-2}$ | 1.175 | 0.086 | 13.611 | <0.001*** |
| $u_{A,t-24}$ | 0.394 | 0.087 | 4.521 | <0.001*** |
| | Std. Dev | | | $R^2$ |
| Random effect | 0.551 | | Marginal | 0.188 |
| Residual | 1.000 | | Conditional | 0.208 |

| Vegetation (V) | | | | |
|---|---|---|---|---|
| | Estimate | Std. Error | z value | p-value |
| (Intercept) | -0.496 | 0.187 | -2.655 | 0.008** |
| *PhasePost-closure* | -0.252 | 0.055 | -4.593 | <0.001*** |
| $h_{FS}$ | -1.870 | 0.443 | -4.223 | <0.001*** |
| $u_{V,t-1}$ | 2.597 | 0.061 | 42.240 | <0.001*** |
| $u_{V,t-2}$ | 0.853 | 0.064 | 13.327 | <0.001*** |
| $u_{V,t-24}$ | 0.382 | 0.061 | 6.256 | <0.001*** |
| | Std. Dev | | | $R^2$ |
| Random effect | 0.361 | | Marginal | 0.298 |
| Residual | 1.000 | | Conditional | 0.314 |

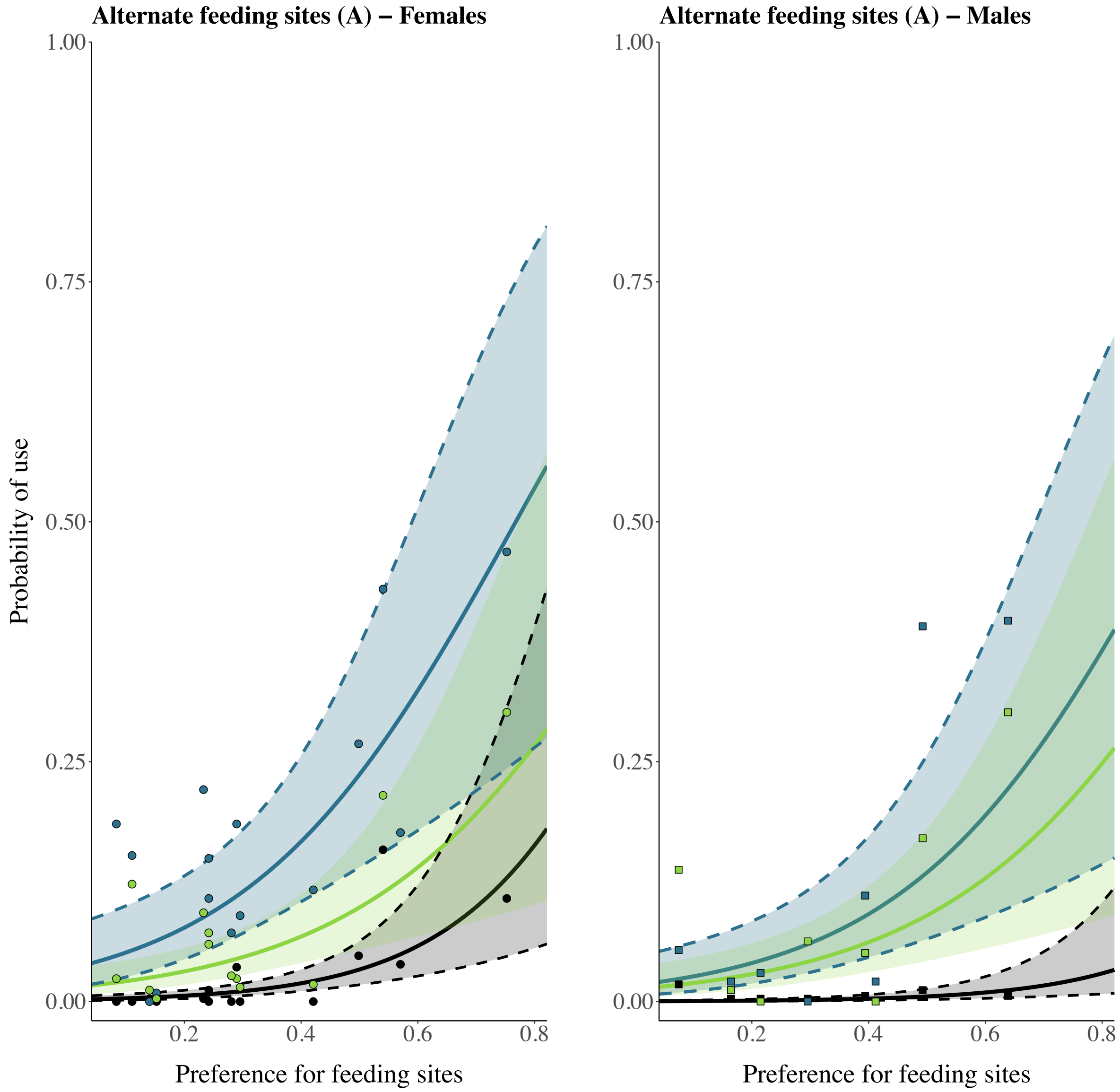


Figure S1. Roe deer shifts in use of alternate feeding sites (A, $u_{A,t}$, y-axis) during the experiment (pre-closure: black; closure: blue and post-closure: green), as a function of preference for feeding sites (y-axis) and *Sex* (left panel: females, right panel: males). Model predictions are plotted as solid lines (95% confidence interval: ribbon) and mean relative use by dots (females) or squares (males). The model predictions do not consider the influence of resource lags at 1, 2 and 24 h.

Table S2. Summary of the full models for the use of the manipulated feeding site ($u_{M,t}$) and vegetation ($u_{V,t}$). The summary for the use of alternate feeding sites ($u_{A,t}$) is not shown as the full model is also the final model (see Supplementary S6: Table S1). The models include experimental phase (*Phase*; reference level: *Pre-closure*), preference for feeding sites ($h_{FS}$), *Sex* (reference level: female, *F*), the interactions of *Phase* with both $h_{FS}$ and *Sex,* and the resource variables at lags 1, 2 and 24 hours (e.g., $u_{M,t-1}$, $u_{M,t-2}$ and $u_{M,t-24}$) as fixed effects, and animal-year as a random intercept. For the vegetation model, the data included only the Closure and Post-closure phases since the average $u_{V,t}$ during pre-closure was used to calculate $h_{FS}$. The reference levels used for *Phase* were *Pre-closure* for $u_{M,t}$, and *Closure* for $u_{V,t}$.

| Manipulated FS (M) | | | | |
|---|---|---|---|---|
| | Estimate | Std. Error | z value | p-value |
| (Intercept) | -3.325 | 0.117 | -28.359 | <0.001*** |
| *PhaseClosure* | -0.715 | 0.183 | -3.915 | <0.001*** |
| *PhasePost-closure* | -0.060 | 0.131 | -0.458 | 0.647 |
| $h_{FS}$ | 1.724 | 0.292 | 5.913 | 0.001** |
| *Sex* | 0.006 | 0.108 | 0.059 | 0.953 |
| *PhaseClosure*:$h_{FS}$ | -1.705 | 0.469 | -3.637 | <0.001*** |
| *PhasePost-closure:*$h_{FS}$ | -0.401 | 0.330 | -1.216 | 0.224 |
| *PhaseClosure*:*SexM* | 0.216 | 0.160 | 1.348 | 0.178 |
| *PhasePost-closure:SexM* | -0.185 | 0.124 | -1.487 | 0.137 |
| $u_{M,t-1}$ | 3.152 | 0.065 | 48.505 | <0.001*** |
| $u_{M,t-2}$ | 0.898 | 0.068 | 13.255 | <0.001*** |

| | | | | |
|---|---|---|---|---|
| $u_{M,t-24}$ | 0.737 | 0.062 | 11.894 | <0.001*** |
| | Std. Dev | | | $R^2$ |
| Random effect | 0.163 | | Marginal | 0.347 |
| Residual | 1.000 | | Conditional | 0.350 |

| Vegetation (V) | | | | |
|---|---|---|---|---|
| | Estimate | Std. Error | z value | p-value |
| (Intercept) | -0.565 | 0.198 | -2.846 | 0.004** |
| *PhasePost-closure* | -0.235 | 0.117 | -2.010 | 0.044* |
| $h_{FS}$ | -1.904 | 0.461 | -4.131 | <0.001*** |
| *Sex* | 0.243 | 0.177 | 1.378 | 0.168 |
| *PhasePost-closure:*$h_{FS}$ | 0.040 | 0.292 | 0.136 | 0.892 |
| *PhasePost-closure:SexM* | -0.091 | 0.116 | -0.785 | 0.432 |
| $u_{V,t-1}$ | 2.596 | 0.061 | 42.222 | <0.001** |
| $u_{V,t-2}$ | 0.852 | 0.064 | 13.311 | <0.001** |
| $u_{V,t-24}$ | 0.381 | 0.061 | 6.240 | <0.001** |
| | Std. Dev | | | $R^2$ |
| Random effect | 0.351 | | Marginal | 0.300 |
| Residual | 1.000 | | Conditional | 0.315 |

Table S3. Summary of the best models for the use of the manipulated feeding site ($u_{M,t}$), alternate feeding sites ($u_{A,t}$) and vegetation ($u_{V,t}$) without considering the resource variables at lags 1, 2 and 24 hours (e.g., $u_{M,t-1}$, $u_{M,t-2}$ and $u_{M,t-24}$). The models include experimental phase (*Phase*), preference for feeding sites ($h_{FS}$), *Sex* (reference level: female, *F*), the interaction of *Phase* with both $h_{FS}$ (for $u_{M,t}$ and $u_{A,t}$) and *Sex* (for $u_{A,t}$, only) as fixed effects, and animal-year as a random intercept. For the vegetation model, the data included only the Closure and Post-closure phases since the average $u_{V,t}$ during pre-closure was used to calculate $h_{FS}$. The reference levels used for *Phase* were *Pre-closure* for $u_{M,t}$ and $u_{A,t}$, and *Closure* for $u_{V,t}$.

| Manipulated FS (M) | | | | |
|---|---|---|---|---|
| | Estimate | Std. Error | z value | p-value |
| (Intercept) | -2.466 | 0.173 | -14.294 | <0.001*** |
| *PhaseClosure* | -0.908 | 0.146 | -6.234 | <0.001*** |
| *PhasePost-closure* | -0.058 | 0.095 | -0.607 | 0.544 |
| $h_{FS}$ | 4.576 | 0.453 | 10.093 | <0.001*** |
| *PhaseClosure*:$h_{FS}$ | -4.269 | 0.382 | -11.182 | <0.001*** |
| *PhasePost-closure:*$h_{FS}$ | -1.639 | 0.234 | -7.002 | <0.001*** |

| | Std. Dev |
|---|---|
| Random effect | 0.370 |
| Residual | 1.000 |

| | $R^2$ |
|---|---|
| Marginal | 0.157 |
| Conditional | 0.174 |

| Alternate FS (A) | | | | |
|---|---|---|---|---|
| | Estimate | Std. Error | z value | p-value |
| (Intercept) | -6.246 | 0.507 | -12.31 | <0.001*** |
| *PhaseClosure* | 2.887 | 0.252 | 11.463 | <0.001*** |

| | | | | |
|---|---|---|---|---|
| *PhasePost-closure* | 2.013 | 0.262 | 7.678 | <0.001*** |
| $h_{FS}$ | 5.762 | 1.192 | 4.833 | <0.001*** |
| *Sex* | -1.862 | 0.519 | -3.59 | <0.001*** |
| *PhaseClosure*:$h_{FS}$ | -1.387 | 0.463 | -2.993 | 0.003** |
| *PhasePost-closure:*$h_{FS}$ | -1.733 | 0.481 | -3.605 | <0.001*** |
| *PhaseClosure*:*SexM* | 1.171 | 0.272 | 4.313 | <0.001*** |
| *PhasePost-closure:SexM* | 1.764 | 0.277 | 6.368 | <0.001*** |

| | Std. Dev | | | $R^2$ |
|---|---|---|---|---|
| Random effect | 1.007 | | Marginal | 0.160 |
| Residual | 1.000 | | Conditional | 0.208 |

| Vegetation (V) | | | | |
|---|---|---|---|---|
| | Estimate | Std. Error | z value | p-value |
| (Intercept) | 2.898 | 0.306 | 9.473 | <0.001*** |
| *PhasePost-closure* | -0.490 | 0.044 | -11.219 | <0.001*** |
| $h_{FS}$ | -3.755 | 0.802 | -4.680 | <0.001*** |

| | Std. Dev | | | $R^2$ |
|---|---|---|---|---|
| Random effect | 0.721 | | Marginal | 0.076 |
| Residual | 1.000 | | Conditional | 0.154 |

Table S4. Summary of the final models for the use of the manipulated feeding site ($u_{M,t}$), alternate feeding sites ($u_{A,t}$) and vegetation ($u_{V,t}$) when incorporating the two outlier animals – F4-2017 and F16-2017 – regarding the availability of A. The models include experimental phase (*Phase*), preference for feeding sites ($h_{FS}$), the interaction of *Phase* with both $h_{FS}$ (for $u_{M,t}$ and $u_{O,t}$) and *Sex* (for $u_{A,t}$, only), and the response variables at lags 1, 2 and 24 hours (e.g., $u_{M,t-1}$, $u_{M,t-2}$ and $u_{M,t-24}$) as fixed effects, and animal-year as a random intercept. For the vegetation model, the data included only the Closure and Post-closure phases since the average $u_{V,t}$ during pre-closure was used to calculate $h_{FS}$. The reference levels used for *Phase* were *Pre-closure* for $u_{M,t}$ and $u_{A,t}$, and *Closure* for $u_{V,t}$.

| Manipulated FS (M) | | | | |
|---|---|---|---|---|
| | Estimate | Std. Error | z value | p-value |
| (Intercept) | -3.342 | 0.122 | -27.432 | <0.001*** |
| *PhaseClosure* | -0.774 | 0.168 | -4.613 | <0.001*** |
| *PhasePost-closure* | -0.133 | 0.125 | -1.062 | 0.288 |
| $h_{FS}$ | 1.676 | 0.306 | 5.483 | <0.001*** |
| *PhaseClosure*:$h_{FS}$ | -1.087 | 0.413 | -2.633 | 0.008** |
| *PhasePost-closure:*$h_{FS}$ | -0.359 | 0.309 | -1.162 | 0.245 |
| $u_{M,t-1}$ | 3.170 | 0.062 | 50.947 | <0.001*** |
| $u_{M,t-2}$ | 0.888 | 0.065 | 13.699 | <0.001*** |
| $u_{M,t-24}$ | 0.757 | 0.059 | 12.817 | <0.001*** |

| | Std. Dev | | | $R^2$ |
|---|---|---|---|---|
| Random effect | 0.202 | | Marginal | 0.355 |

| | | | | |
|---|---|---|---|---|
| Residual | 1.000 | | Conditional | 0.359 |

| Alternate FS (A) | | | | |
|---|---|---|---|---|
| | Estimate | Std. Error | z value | p-value |
| (Intercept) | -5.504 | 0.429 | -12.824 | <0.001*** |
| *PhaseClosure* | 1.741 | 0.251 | 6.933 | <0.001*** |
| *PhasePost-closure* | 1.234 | 0.262 | 4.711 | <0.001*** |
| $h_{FS}$ | 2.794 | 0.989 | 2.825 | 0.005** |
| *Sex* | -1.078 | 0.443 | -2.433 | 0.015* |
| *PhaseClosure*:$h_{FS}$ | -1.416 | 0.513 | -2.760 | 0.006** |
| *PhasePost-closure:*$h_{FS}$ | -1.484 | 0.536 | -2.766 | 0.006** |
| *PhaseClosure*:*SexM* | 0.837 | 0.283 | 2.953 | 0.003** |
| *PhasePost-closure:SexM* | 1.124 | 0.291 | 3.856 | <0.001*** |
| $u_{A,t-1}$ | 3.065 | 0.077 | 39.777 | <0.001*** |
| $u_{A,t-2}$ | 1.036 | 0.082 | 12.646 | <0.001*** |
| $u_{A,t-24}$ | 0.477 | 0.080 | 5.982 | <0.001*** |

| | Std. Dev | | | $R^2$ |
|---|---|---|---|---|
| Random effect | 0.809 | | Marginal | 0.174 |
| Residual | 1.000 | | Conditional | 0.220 |

| Vegetation (V) | | | | |
|---|---|---|---|---|
| | Estimate | Std. Error | z value | p-value |
| (Intercept) | -0.584 | 0.172 | -3.391 | 0.001** |

| | | | | |
|---|---|---|---|---|
| *PhasePost-closure* | -0.258 | 0.052 | -4.974 | 0.001** |
| $h_{FS}$ | -1.772 | 0.391 | -4.536 | 0.001** |
| $u_{V,t-1}$ | 2.647 | 0.059 | 45.039 | <0.001*** |
| $u_{V,t-2}$ | 0.793 | 0.061 | 12.979 | <0.001*** |
| $u_{V,t-24}$ | 0.473 | 0.057 | 8.294 | <0.001*** |
| | -0.584 | 0.172 | -3.391 | |

| | Std. Dev | | | $R^2$ |
|---|---|---|---|---|
| Random effect | 0.338 | | Marginal | 0.310 |
| Residual | 1.000 | | Conditional | 0.324 |

**Supplementary S7: Sensitivity analysis**

We evaluated the sensitivity of our model outputs to the choice of buffer size, used to evaluate feeding site attendance. We tested six buffer sizes calculated as a function of mean roe deer step length, $l$ (61.2 meters): $l$ multiplied by 0.5, 1, 1.5, 2, 3 and 4 (i.e., 30.6, 61.2, 91.8, 122.4, 183.6 and 244.8 m, respectively).

The parameter estimates for both the largest and smallest buffer sizes are characterized by large confidence intervals: $0.5l$ for preference for feeding sites ($h_{FS}$) and its interaction with experimental phase ($Phase$:$h_{FS}$), and $4l$ for the model intercepts and experimental phase ($Phase$). The estimates for $0.5l$ deviates from all other buffer sizes. The estimates and associated confidence intervals of the intermediary, most meaningful buffer sizes – $1l$, $1.5l$ and $2l$ – are consistent for all developed models. The two outlier animals – F4-2017 and F16-2017 – were not included in the comparisons for the resource use models ($u_{M,t}$, $u_{A,t}$ and $u_{V,t}$).

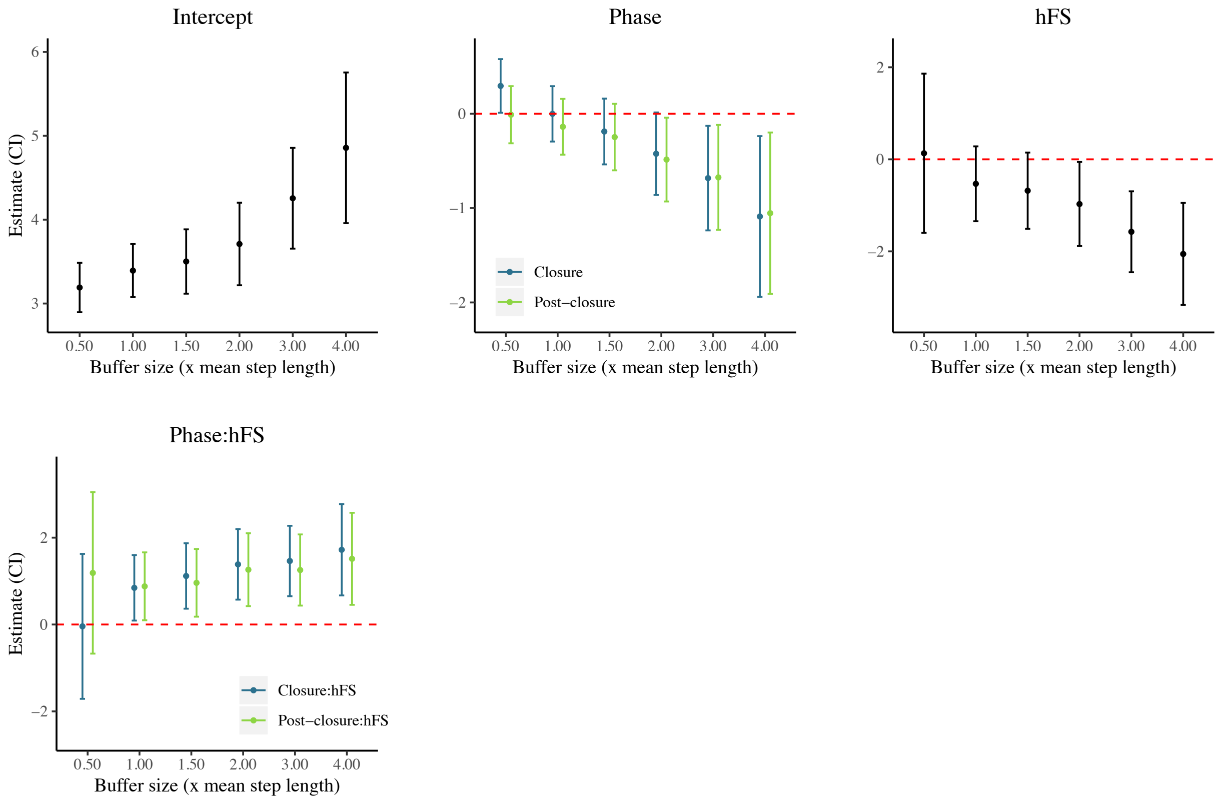


Figure S1. Sensitivity of the home range size (95%UD) model to the choice of buffer size (x-axis) used to define feeding site (FS) attendance. The estimates include the intercept, experimental phase (*Phase*), preference for FS ($h_{FS}$) and their interaction ($Phase$:$h_{FS}$). Buffer size is expressed as a multiple (0.5, 1.0, 1.5, 2.0, 3.0 or 4.0) of the mean roe deer step length, $l$ (i.e., 61.2 m).

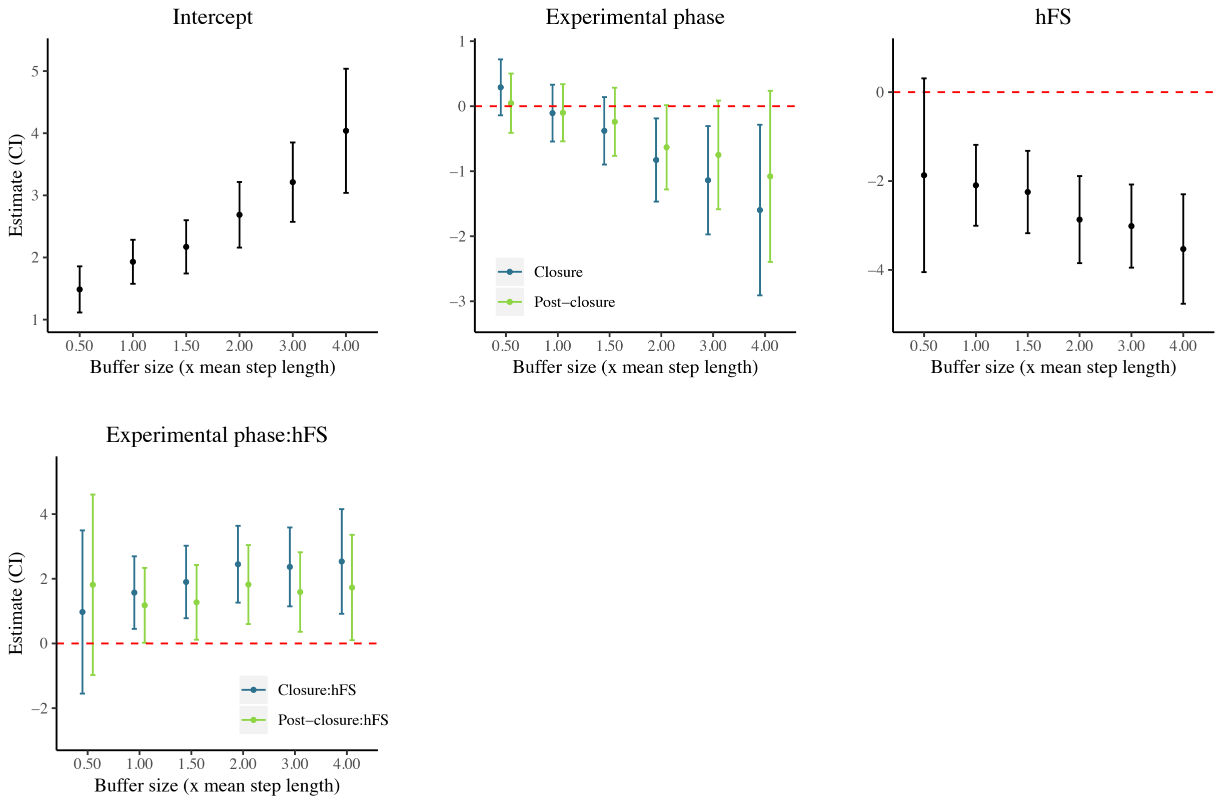


Figure S2. Sensitivity of the core area size (50% UD) model to the choice of buffer size (x-axis) used to define feeding site (FS) attendance. The estimates include the intercept, experimental phase (*Phase*), preference for FS ($h_{FS}$) and their interaction (*Phase*:$h_{FS}$). Buffer size is expressed as a multiple (0.5, 1.0, 1.5, 2.0, 3.0 or 4.0) of the mean roe deer step length, $l$ (i.e., 61.2 m).

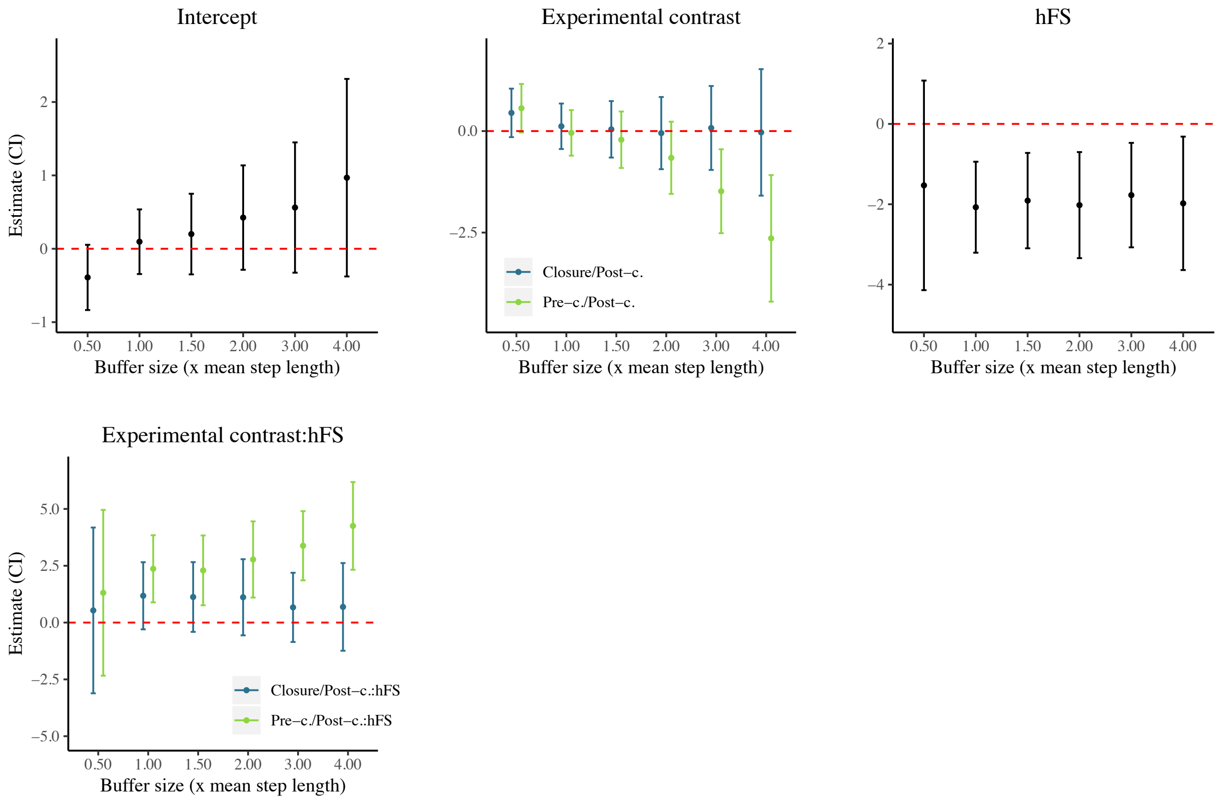


Figure S3. Sensitivity of the space-use overlap model to the choice of buffer size (x-axis) used to define feeding site (FS) attendance. The estimates include the intercept, experimental phase (*Phase*), preference for FS ($h_{FS}$) and their interaction (*Phase*:$h_{FS}$). Buffer size is expressed as a multiple (0.5, 1.0, 1.5, 2.0, 3.0 or 4.0) of the mean roe deer step length, $l$ (i.e., 61.2 m).

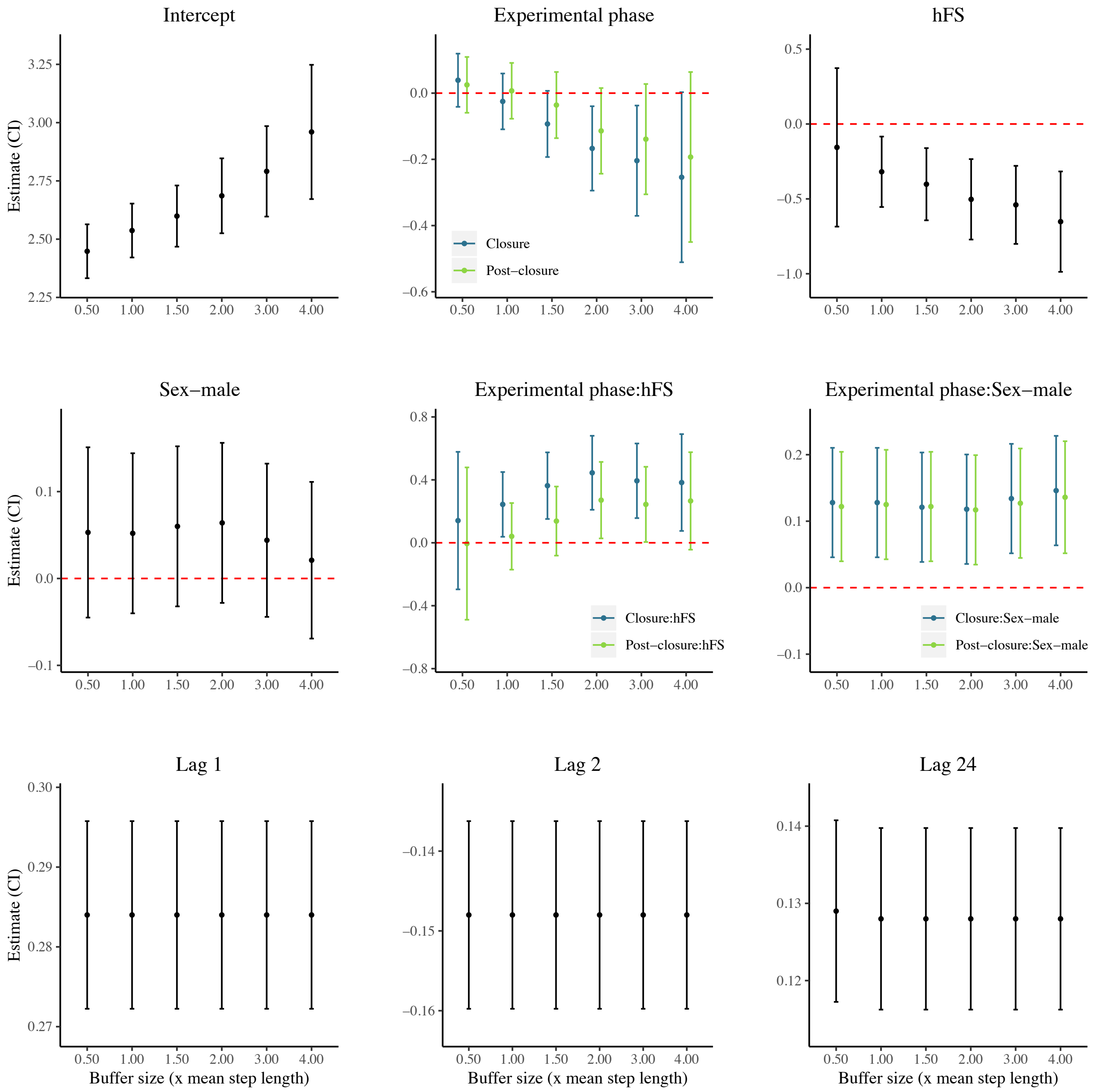


Figure S4. Sensitivity of the step length ($s_t$) model to the choice of buffer size (x-axis) used to define feeding site (FS) attendance. The estimates include the intercept, experimental phase (*Phase*), preference for FS ($h_{FS}$), *Sex*, the interactions of *Phase* with both $h_{FS}$ and *Sex*, and the step length at lags 1, 2 and 24 hours ($s_{t-1}$, $s_{t-2}$ and $s_{t-24}$). Buffer size is expressed as a multiple (0.5, 1.0, 1.5, 2.0, 3.0 or 4.0) of the mean roe deer step length, $l$ (i.e., 61.2 m).

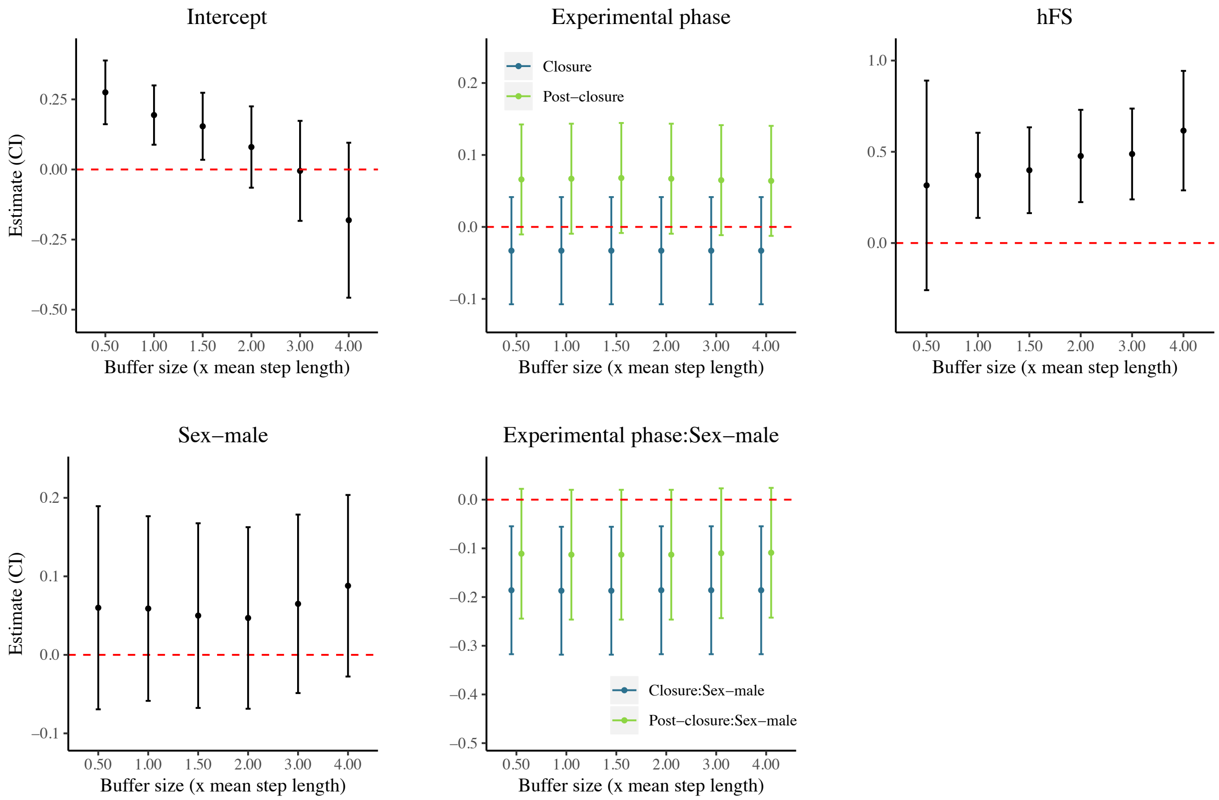


Figure S5. Sensitivity of the absolute turn angle model ($\varphi_t$) model to the choice of buffer size (x-axis) used to define feeding site (FS) attendance. The estimates include the intercept, experimental phase (*Phase*), preference for FS ($h_{FS}$), *Sex*, and the interaction of *Phase Sex*. Buffer size is expressed as a multiple (0.5, 1.0, 1.5, 2.0, 3.0 or 4.0) of the mean roe deer step length, $l$ (i.e., 61.2 m).

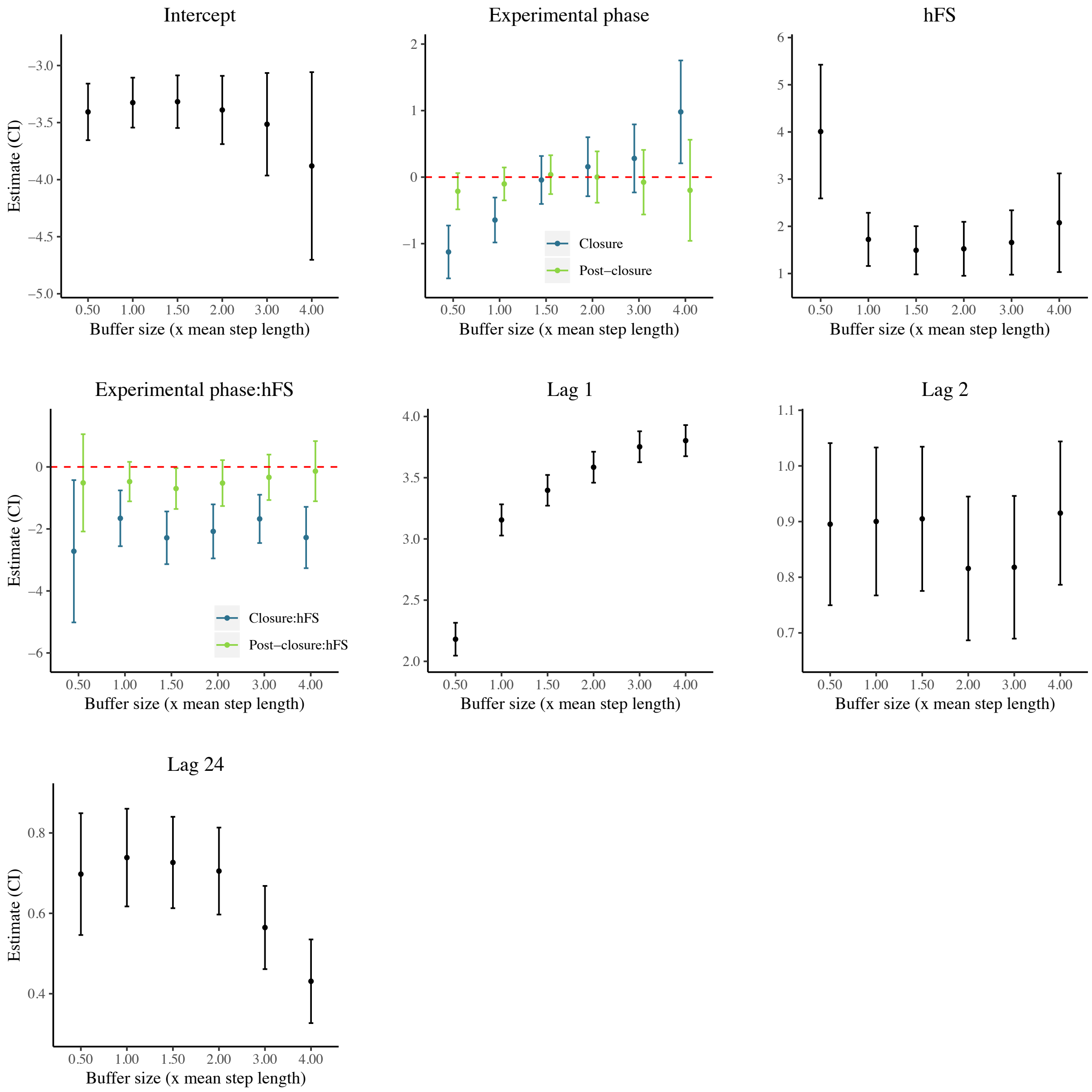


Figure S6. Sensitivity of the manipulated feeding site use ($u_{M,t}$) model to the choice of buffer size (x-axis) used to define feeding site (FS) attendance. The estimates include the intercept, experimental phase (*Phase*), preference for FS ($h_{FS}$), the interaction of *Phase* with $h_{FS}$, and the use of M at lags 1, 2 and 24 hours ($u_{M,t-1}$, $u_{M,t-2}$ and $u_{M,t-24}$). Buffer size is expressed as a multiple (0.5, 1.0, 1.5, 2.0, 3.0 or 4.0) of the mean roe deer step length, $l$ (i.e., 61.2 m).

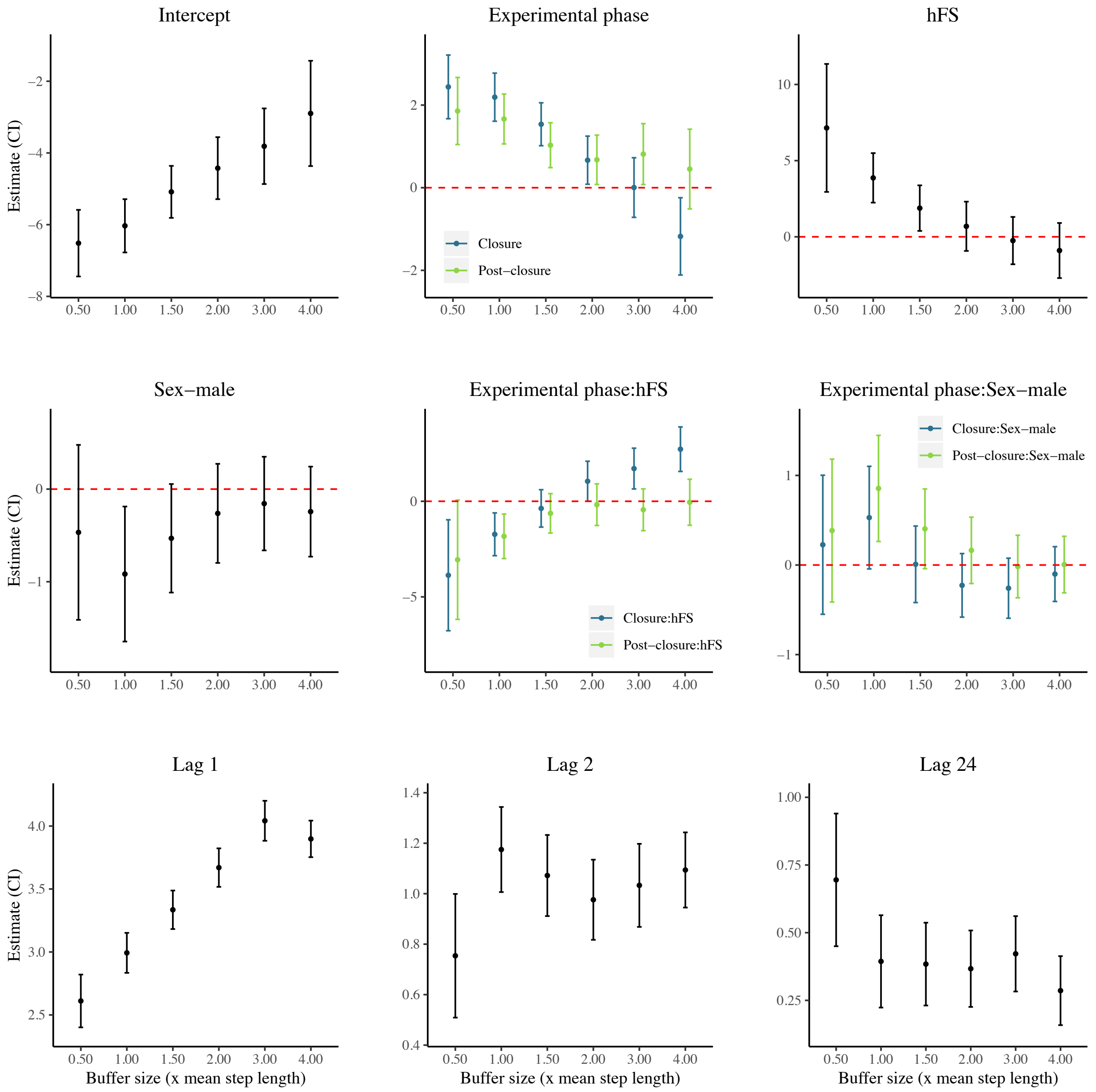


Figure S7. Sensitivity of the alternate feeding site use ($u_{A,t}$) model to the choice of buffer size (x-axis) used to define feeding site (FS) attendance. The estimates include the intercept, experimental phase (*Phase*), preference for FS ($h_{FS}$), *Sex*, the interactions of *Phase* with both $h_{FS}$ and *Sex*, and the use of A at lags 1, 2 and 24 hours ($u_{A,t-1}$, $u_{A,t-2}$ and $u_{A,t-24}$). Buffer size is expressed as a multiple (0.5, 1.0, 1.5, 2.0, 3.0 or 4.0) of the mean roe deer step length, $l$ (i.e., 61.2 m).

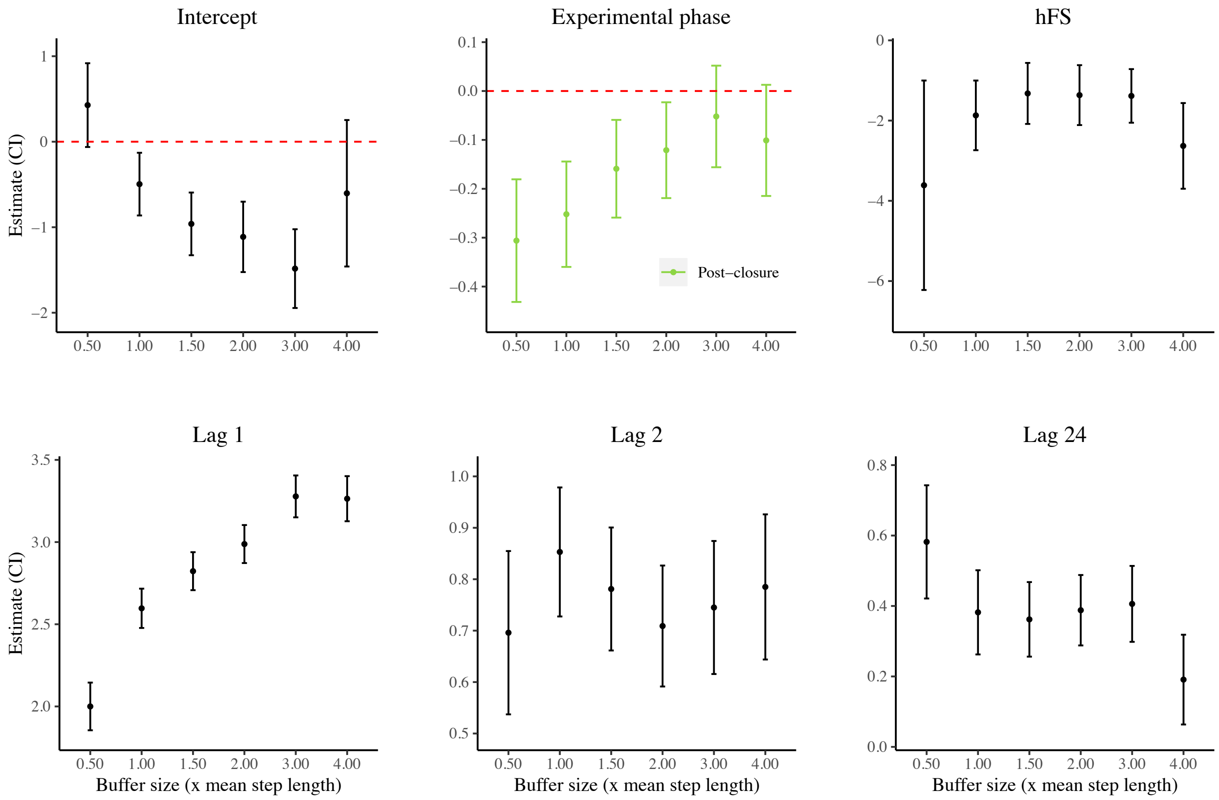


Figure S8. Sensitivity of the vegetation use ($u_{V,t}$) model to the choice of buffer size (x-axis) used to define feeding site (FS) attendance. The estimates include the intercept, experimental phase (*Phase*), preference for FS ($h_{FS}$), and the use of V at lags 1, 2 and 24 hours ($u_{V,t-1}$, $u_{V,t-2}$ and $u_{V,t-24}$). Buffer size is expressed as a multiple (0.5, 1.0, 1.5, 2.0, 3.0 or 4.0) of the mean roe deer step length, $l$ (i.e., 61.2 m).

**Supplementary S8: Personality correlates of preference for feeding sites**

Following Bonnot et al.[1], we estimated individual boldness, an established personality trait[2], using two indexes: the body temperature at capture (a known physiological parameter of individual stress [3]), and a behavioural score of individual reactivity during the capture ('boldness' index). We evaluated boldness for each animal-year (see Supplementary S3: Table S3.1). We could not assess the repeatability of these indexes because of the scarce number of recaptured individuals across the three sampling years (n=5). However, both metrics have already been shown to be estimates of individual stress and personality in roe deer (*Capreolus capreolus*) with a moderate to high degree of repeatability[1].

We measured the body (rectal) temperature during capture while handling and marking the roe deer. As for the behavioural score, we readapted the behavioural index described in Bonnot et al.[1] to the capture methodology used in our study area. We computed the 'boldness' index as the sum of two behavioural scores estimated at capture i.e., the reactivity during handling (ranging from 0 to 4, see Table S1) and the flight behaviour at the release (ranging from 0 to 4, see Table S1). The boldness index ranged from 0 to 8, where 0 denotes a 'bold' individual and 8 denotes a 'shy' (very reactive) individual at capture.

Measurements of body temperature and behavioural score at capture were available for 22 and 24 animal-years, respectively (out of 25). We found that the body temperature and the boldness index were significantly correlated ($r = 0.51$, $p = 0.021$). Individual preference for FS was marginally correlated with body temperature ($r = - 0.37$, $p = 0.084$) but not with the boldness index ($r = - 0.23$, $p = 0.29$).

This analysis suggests a correlation between roe deer personality, and in particular individual boldness, and the preference for feeding sites. We argue that the marginal significance that we found is likely to be explained by the relatively small sample size available for this analysis.

Table S1. Description of the handling and release behaviour scores.

| Value | Handling behaviour | Release behaviour |
|---|---|---|
| 0 | Calm. No resistance. No kicking with legs. No barking. | The animal goes away slowly. It stops to look back several times. |
| 1 | Calm. Almost no kicking. Only a couple of barking. | The animal runs away but it stops after a short distance. |
| 2 | Kicking and barking some time but alternating calm phases. | The animal runs away, never stopping until when it is out of the field view. |
| 3 | Stressed. Kicking and barking but it can be managed. | The animal fells over and jumps attempting to remove the collar and to get free from the capture team. |
| 4 | Very stressed. Very hard to handle. Impossible to take biometric measurements in a proper way. | The animal lies on the ground. It is unable to stand up by itself. |
